# BIT: Bayesian Identification of Transcriptional Regulators from Epigenomics-Based Query Region Sets

**DOI:** 10.1101/2024.06.02.597061

**Authors:** Zeyu Lu, Lin Xu, Xinlei Wang

## Abstract

Transcriptional regulators (TRs) are master controllers of gene expression and play a critical role in both normal tissue development and disease progression. However, existing computational methods for identification of TRs regulating specific biological processes have significant limitations, such as relying on distance on a linear chromosome or binding motifs that have low specificity. Many also use statistical tests in ways that lack interpretability and rigorous confidence measures. We introduce BIT, a novel Bayesian hierarchical model for *in-silico* TR identification. Leveraging a comprehensive library of TR ChIP-seq data, BIT offers a fully integrated Bayesian approach to assess genome-wide consistency between user-provided epigenomic profiling data and the TR binding library, enabling the identification of critical TRs while quantifying uncertainty. It avoids estimation and inference in a sequential manner or numerous isolated statistical tests, thereby enhancing accuracy and interpretability. BIT successfully identified critical TRs in perturbation experiments, functionally essential TRs in various cancer types, and cell-type-specific TRs within heterogeneous cell populations, offering deeper biological insights into transcriptional regulation.

## Introduction

Transcriptional regulators (TRs), comprising transcription factors, cofactors, chromatin modulators, and various regulatory molecules, play a key role in regulating gene transcription^1-3^. TRs are essential in diverse cellular processes, including growth, differentiation, morphogenesis, and apoptosis^4-6^. Dysfunction of TRs can lead to various diseases such as cancers^7-9^, diabetes^10,11^, and heart diseases^12,13^. Thus, gaining insights into the identity and function of TRs that regulate specific biological processes is essential in a broad range of biological and medical research, particularly in finding new biomarkers and therapeutic targets^14-16^. Because traditional wet lab experiments to identify TRs are labor-intensive and time-consuming, researchers are increasingly turning to efficient statistical and machine learning methods for TR identification, especially those leveraging next-generation sequencing (NGS) data.

The *in-silico* identification of TRs regulating a specific biological process typically requires two key components. The first component is input data that are often generated from cost-effective techniques, such as a set of genes identified from transcriptomics techniques (e.g., microarray, RNA-seq) or a set of accessible chromatin regions derived from epigenomic profiling techniques^17^ (e.g., ATAC-seq). These Input data typically reflect the downstream effects of key TR activity or provide epigenomic features closely associated with TR binding sites. For example, differential gene expression between diseased and normal samples is often attributed to the change in key TR activity^18-20^.

In our recent comprehensive review^21^, we discussed the limitations of using gene sets as input. This approach requires assigning each gene a representative chromatin location, typically the transcription start site (TSS). The association between a TR and a gene is then assessed based on the linear distance between the binding sites of the TR and the TSS of this gene, which often underestimates long-range interactions, such as distant enhancers that locate thousands of base pairs away from their target genes^22^. In contrast, alternative input such as chromatin accessibility profiling data directly correlate with TR binding activity^23-25^. As accessible chromatin is a prerequisite for TR binding, functional TRs can be inferred as those with binding sites enriched in the input accessible chromatin regions, thereby avoiding the use of linear distance. In addition, epigenomic profiling techniques such as ATAC-seq have gained increasing popularity over the past decade. The abundance of such datasets has stimulated the development of computational methods to uncover new insights into transcriptional regulation.

The second component of *in-silico* TR identification is a reference library consisting of TR ChIP-seq data or TR binding motifs. TRs bind to specific cis-regulatory elements to exert their regulatory function^2^, and thus, data that accurately capture TR binding sites serve as a valuable reference. TR ChIP-seq provides an accurate representation of in-vivo TR binding profile^26^. However, each ChIP-seq dataset can only decipher one TR’s binding profile under a specific condition. It is not feasible to conduct thousands of ChIP-seq experiments for one single condition to measure every TR’s binding profile. Therefore, researchers often rely on computational methods to infer functional TRs based on a library of binding profiles collected from thousands of existing ChIP-seq experiments that cover many known TRs. In addition, TR binding sites can also be inferred by TR binding motifs, which are DNA sequence patterns that reveal TR binding preference.

Despite tremendous successes in advancing biomedical research, existing computational methods can suffer from various limitations, regardless of the reference type. (i) For methods (e.g., WhichTF^27^ and HOMER^28^) that leverage TR binding motifs as reference, the highly similar motifs shared among many TRs make accurate identification challenging^29^. In addition, motifs alone cannot characterize any context-specific binding pattern, and the lack of well-defined motifs for some TRs further reduces the reliability of using motifs for TR identification^2,30^. (ii) For methods using TR ChIP-seq data (e.g., BART^31^ and ChIP-Atlas^32^), each TR’s binding profile can vary in different cell types or environmental conditions. However, published methods have no rigorous treatment of the within-TR heterogeneity when dealing with multiple binding profiles from the same TR. Further, TRs with more datasets can benefit from randomness and have a greater chance of being ranked higher in the final output, leading to biased results^21^. (iii) Most existing methods use frequentist approaches for measuring TR’s statistical significance, lacking coherent probabilistic interpretation or uncertainty estimation. Given the dynamic nature of TR binding activity, the ability to quantify uncertainty is not just beneficial but essential.

To address these limitations, we propose a novel Bayesian hierarchical model for TR identification, named BIT (**B**ayesian **I**dentification of **T**ranscriptional regulators from epigenomics-based query regions sets). The input of BIT is a set of user-provided epigenomic regions containing the chromosome numbers and coordinates of the regions, typically referred to as “peaks”, often derived from genome-wide epigenomic profiling of specific biological processes. Compared to existing methods, BIT offers the following advantages.

First, BIT leverages over 10,000 TR ChIP-seq datasets gathered from previous studies covering a large collection of TRs. This approach bypasses the need of computationally predicted motifs, which can lead to unreliable TR predictions in motif-based methods^28,33,34^. In addition, the context-specific binding patterns captured by TR ChIP-seq will not be neglected.

Second, through a unique hierarchical model setup, BIT integrates information across multiple TRs and across multiple ChIP-seq datasets of the same TRs, appropriately accounting for both between- and within-TR heterogeneity in binding profiles. This formal model-based approach enables BIT to estimate the uncertainty associated with varying context-specific binding patterns existing within different ChIP-seq datasets for the same TR, leading to a more reliable and robust prediction, as shown in the analysis results of this study. By contrast, many existing methods need to conduct thousands of separate *ad-hoc* statistical tests (e.g., BART^31^ and ChIP-Atlas^32^), which often give higher rankings to TRs with more ChIP-seq datasets^21^.

Third, BIT employs a Bayesian framework to handle its modeling and computational needs. Besides information pooling for improved prediction, a Bayesian approach is inherently attractive because (i) it offers the advantage of readily quantifying the estimation uncertainty, such as providing 95% credible intervals for any quantity of interest (e.g., an estimated parameter or predicted importance score); (ii) in situations when meaningful prior knowledge is available, it enables researchers to formally incorporate such knowledge through prior specification for improved results. In all other situations, with the default non-informative prior setting, it becomes a fully automated procedure that does not require any tuning parameter. In contrast, none of the existing methods can offer all these advantages altogether. The R package of BIT is available on GitHub (https://github.com/ZeyuL01/BIT) along with a detailed manual. We also provide an easy- to-use online portal (http://43.135.174.109:8080/).

## Results

The integration of epigenomic profiling data has proven to be a feasible approach for identifying TRs in biomedical research^24,35,36^. It is well recognized that the binding sites of TRs critical to a specific biological process are primarily enriched in active regulatory regions of downstream genes that drive the process^37^. These epigenomic regions can be profiled by sequencing techniques^17^ (e.g., ATAC-seq), which offer a ‘snapshot’ of genome-wide TR activity, akin to distinct ‘fingerprints’ at a crime scene, providing the clues about individuals involved. To link this TR fingerprints snapshot to specific TRs, we leverage TR ChIP-seq data. In humans and mice, there are approximately 1600 known TRs regulating a wide variety of biological processes, and tens of thousands of TR ChIP-seq experiments have been conducted, each creating a unique reference “fingerprint” for an individual TR (under some specific condition). To use this wealth of data, we pre-processed a substantial collection of TR ChIP-seq datasets, creating a comprehensive library of TR binding profiles (distinctive “fingerprints” for individual TRs) in human and mouse genomes, respectively. We utilized 10,140 human TR ChIP-seq datasets covering 988 TRs, and 5,681 mouse TR ChIP-seq datasets covering 607 TRs. These datasets were sourced from GTRD^38^, one of the most comprehensive databases in epigenomic research (**Fig. S1**). Each library serves as a powerful reference. Users can compare their own snapshots of TR “fingerprints” (from ATAC-seq, etc.) with those in the corresponding library to identify TRs whose binding patterns show the highest consistency, indicating their functional relevance in the biological context under study.

BIT operates based on two key biological principles: (1) If a TR is truly involved in regulating a given biological process, its binding pattern across the genome should closely align with the user-provided input (active regulatory regions) more closely than a TR that is irrelevant. A stronger correlation between TR binding sites and these input regions suggests a greater likelihood that the TR regulates key genes, thus significantly impacting the biological process under investigation. (2) Each TR possesses a distinct binding pattern, enabling its identification even in the presence of some variation. This means that the binding patterns of a specific TR will be more similar across different conditions, compared to the binding patterns of other TRs (**Fig. S2**). This inherent specificity allows for accurate recognition and differentiation of TRs within complex biological systems.

These principles, also used in other bioinformatics tools^27,31,39^, underscore that an active TR involved in a cellular process will exhibit frequent DNA interactions, leading to captured epigenomic regions near its binding sites. In addition, while variations exist in ChIP-seq data for the same TR due to factors like measurement errors, data quality, tissue specificity, and cell-type specificity, the overall consistency stemming from TR binding preferences allows for unique binding characteristics to emerge. This uniqueness, when compared to input epigenomic data, enables the differentiation of functional TRs from non-functional ones.

BIT only requires a set of epigenomic regions containing the chromosome numbers and coordinates of the regions as input (e.g., Peaks called by MACS2 with the common format such as bed/narrowPeak) and consists of two main steps: (i) data pre-processing and (ii) Bayesian computation and posterior inference with a Gibbs sampler (see **Methods** and **Supplementary Note 1**). First, BIT transforms the coordinates of processed peak files to fixed-length binary vectors with 0s and 1s. Second, BIT formulates a Bayesian hierarchical model to integrate user-provided information with TR binding information from the reference library and then uses Markov Chain Monte Carlo (MCMC) to sample from the joint posterior distribution. Finally, BIT assigns each TR a “BIT score” based on posterior draws (**Fig. 1**). Based on the assumptions stated above, the BIT score can be used as the overall importance measure of the TR being functional. Alternatively, it can be interpreted as the Jaccard index that quantifies the average similarity between the binary vectors of the TR’s binding profiles and the binary vector of the input epigenomic regions.

**Figure 1.**
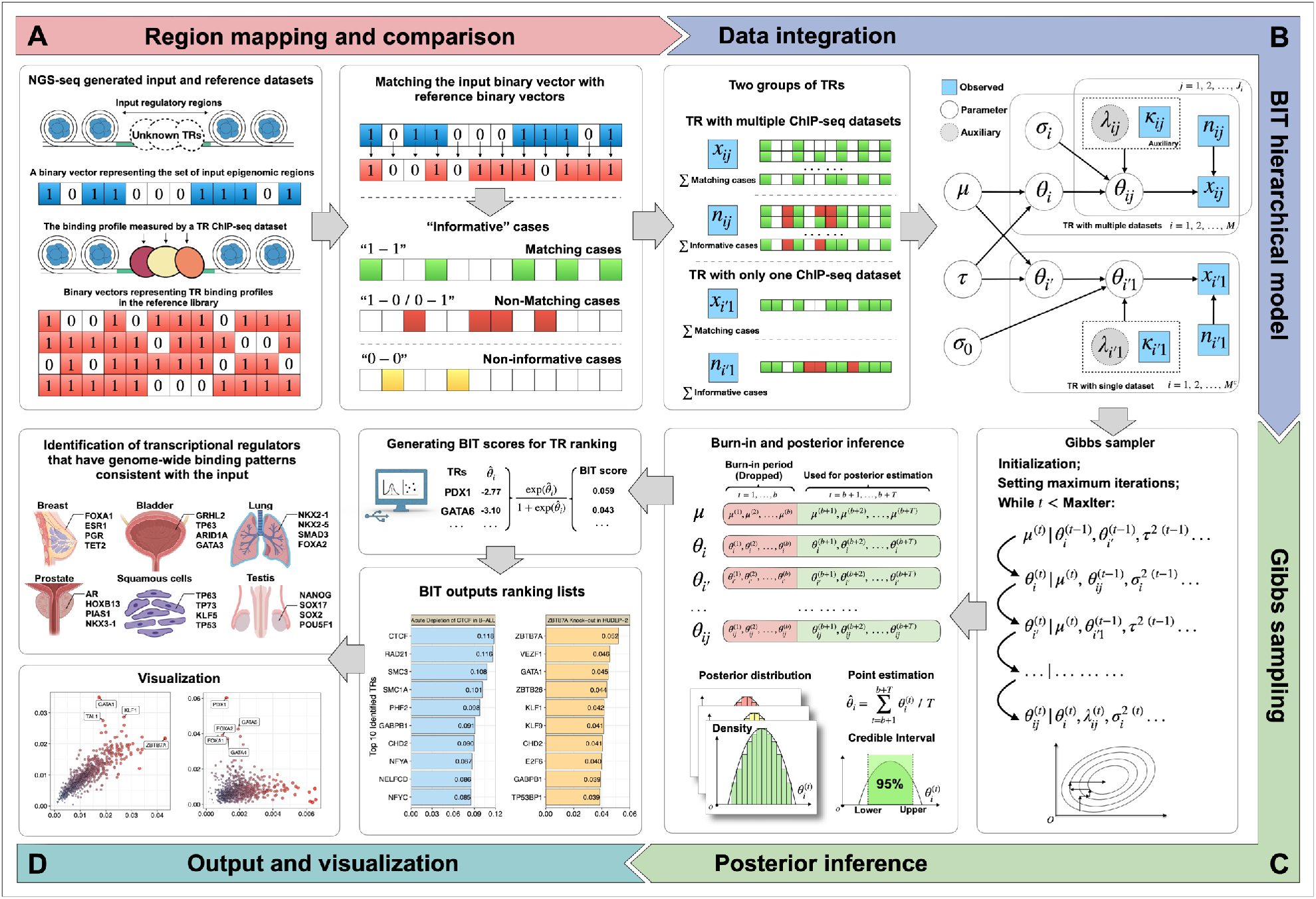
Overview of BIT’s framework. **A**. Transcription regulators (TRs) such as transcription factors, cofactors, and chromatin modulators mostly bind to accessible chromatin regions to regulate gene transcription. NGS techniques such as TR ChIP-seq and ATAC-seq can acquire information on TR binding sites and accessible regions, respectively. Input regions and collected TR ChIP-seq peaks are transformed into binary vectors, where the proportion of the “matching” cases among all “informative” cases serves as the response variable. **B**. BIT is a Bayesian hierarchical model, combined with Pólya-Gamma data augmentation, to integrate information from multiple TRs and multiple ChIP-seq datasets. **C**. A Gibbs sampler is used to draw posterior samples and estimate key model parameters. **D**. TR-level BIT scores are generated and ranked to output a TR ranking list.

To verify BIT’s performance, we first conducted simulations to examine the efficiency of its posterior estimation. As BIT is built on a unique hierarchical model with meaningful parameters, we used a “naïve” method as the baseline for comparison, which estimates the parameters based on straightforward summary statistics as described in **Methods**. Second, we examined BIT’s performance with three different applications: (i) We identified TRs using differentially accessible regions (DARs) derived from TR perturbation experiments. Perturbed TRs should be responsible for the DARs, and therefore these DARs can serve as input with known ground truth. (ii) We tested BIT in TR identification using cancer-type specific accessible epigenomic regions. The identified TRs were cross-validated by findings in previous publications and results derived from CRISPR/Cas9 loss-of-function analysis. (iii) Given the rapid development of single-cell techniques, cell-type-specific accessible regions can now be derived from scATAC-seq data. We further applied BIT to two sets of cell-type-specific accessible regions and validated the BIT predictions. We also benchmarked results from BIT against other state-of-the-art methods in the three applications.

### BIT can estimate model parameters and TR rankings with high accuracy

We explored the performance of BIT through extensive simulations. As will be detailed in the **Methods** section, our BIT model separates candidate TRs in the reference library into two distinct groups: (1) TRs with binding profiles generated from multiple TR ChIP-seq datasets and (2) TRs with only one binding profile generated from a single TR ChIP-seq dataset. We use *i* and *i*′ to index TRs in the first and second groups, respectively. To cover potential scenarios, we first varied the global mean and variance parameters *μ, τ*^2^ to control the distribution of *θ*_*i*_. The *θ*_*i*_ denotes the mean (logit-scale) proportion of matching between the input binary vector representing the user-provided set of epigenomic regions and those representing binding profiles of the *i*th TR in the reference library, measuring its importance relevant to the biological process under study. We also varied 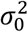, the common variance of *θ*_*i*,1_ within the second TR group, where *θ*_*i*,1_ is the logit-scale proportion of matching between the input binary vector and the reference vector representing the single binding profile of the *i*′th TR. In our simulation, we examined the effect of one such parameter at a time while keeping the other two fixed at their default values (*μ* = −4, *τ*^2^ = 0.75, and 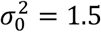), determined based on an exploratory data analysis of TR perturbation experiments in the **Results** Section. The range of the varied parameters was also decided based on the exploratory analysis to cover various real scenarios (*μ* ∈ {−5, −4.5, …, −2.5}, *τ*^2^ ∈ {0.5,0.7, …, 1.5}, and 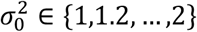.Other parameters were then generated from Gaussian distributions with pre-determined global parameters according to our BIT model. Finally, the number of input “matching” and “informative” bins were generated from a binomial distribution with logit^-1^ transformed *θ*_*ij*_ (*θ*_*i*′1_). We repeated the above steps and generated 100 replicate datasets for each combination of *μ, τ*^2^, and 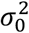.

We further investigated the effect of adjusting the number of candidate TRs (*I*) included in our analysis. Our human reference library covers 988 TRs, and our mouse reference library covers 607 TRs. Both libraries are expected to rapidly grow as more and more TR ChIP-seq datasets are being generated. Thus, we considered *I* = 500, 1000, 1500 in our simulation. In addition, the count of TR ChIP-seq datasets for the *i*th TR was simulated randomly from a distribution fitted using the actual counts within the first group in the human reference library.

We evaluated BIT in three aspects: (i) the mean squared errors (MSEs) for estimating global parameters 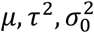 and TR-level importance parameters (*θ*_*i*_’s and *θ*_*i* ′_’s); (ii) the ability to recover the original ranks of TRs according to their importance, measured by Spearman’s rho. (iii) the robustness of BIT to the normality assumption of the BIT model. Comparison was made with a baseline method based on naïve estimation, as detailed in the **Methods** section.

Overall, BIT estimates all global parameters with high efficiency (**Fig. 2A**). First, it is noted that the MSEs of BIT for estimating *μ, τ*^2^, and 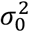 are consistently lower than that of the baseline method and are close to zero in all cases. Second, a lower number of candidate TRs (i.e., smaller *I*) can result in relatively higher MSE, but the difference is much less obvious for BIT. Third, the MSE for the estimation of *τ*^2^ tends to increase as the value of *τ*^2^ goes up. This is reasonable as the increased variability would lead to greater estimation uncertainty. Nevertheless, BIT still significantly outperforms the baseline method in estimating *τ*^2^.

**Figure 2.**
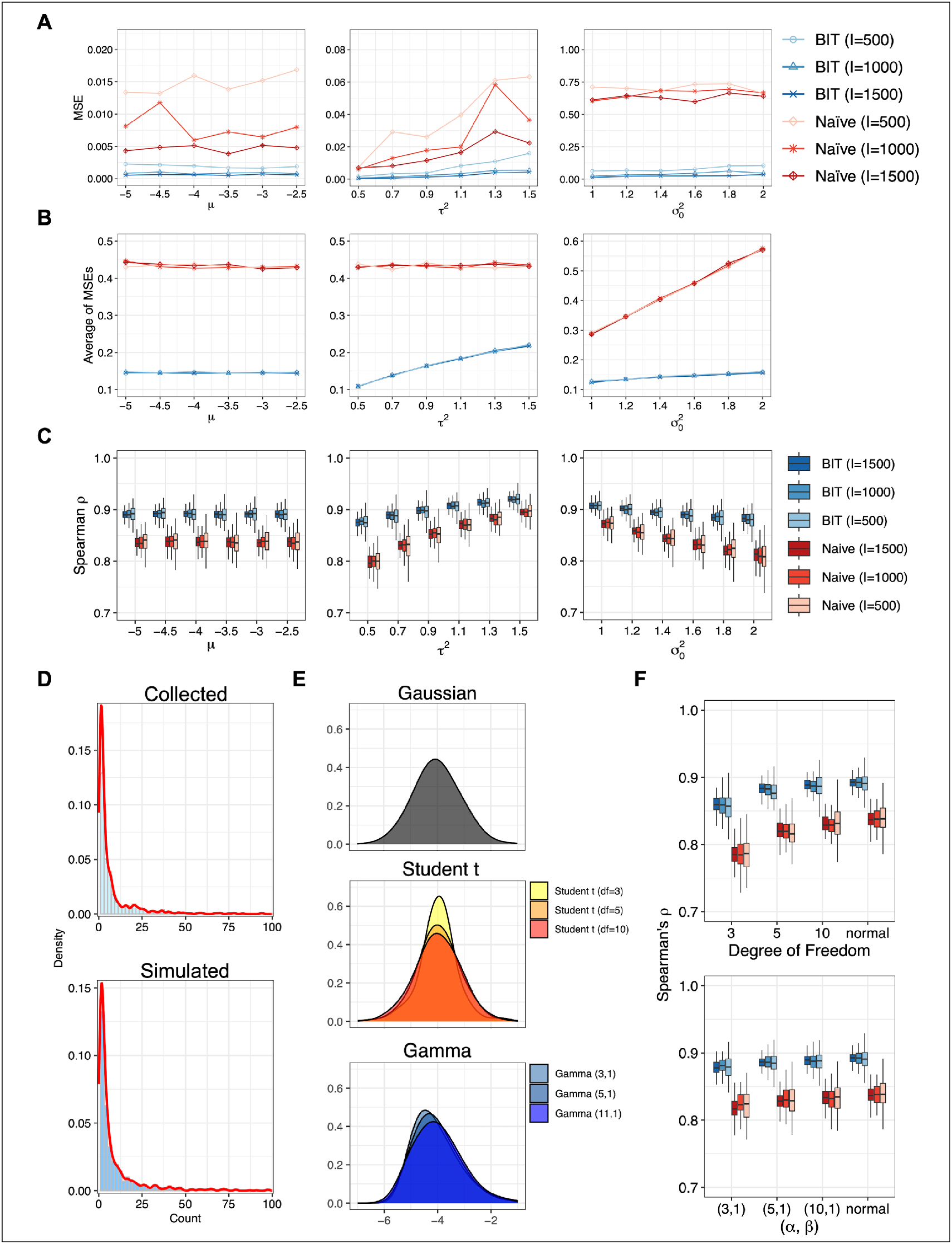
Simulation results under different parameter settings. **A**. Mean squared errors for estimation of global parameters using 100 replicate datasets in each simulation setting. **B**. The average of MSEs for estimation of importance parameters (averaged over different TRs) over 100 replicates. **C**. Boxplots of Spearman’s rho correlation over 100 replicates between ranks from BIT or the baseline method and the true ranks of simulated TRs. **D**. The distributions of simulated vs. real numbers of ChIP-seq datasets of individual TRs. **E**. Data generated in our sensitivity analysis from t and gamma distributions, to examine whether heavy-tailedness and skewness would significantly affect the performance of BIT. **F**. Boxplots of Spearman correlation over 100 replicates under t and gamma distributions, with baseline under normal distribution.

BIT offers improved performance in estimating TR-level importance parameters as well, as evidenced by much lower average MSEs (averaged over the different TRs) in various settings (**Fig. 2B**). Accurate estimation of these parameters is crucial for determining precise TR rankings. We examined Spearman’s rho correlation, which is the Pearson correlation between the rank values of the estimated and true importance parameters of the candidate TRs. We find that for both methods, larger *τ*^2^ yields higher correlation while larger 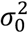 does the opposite (**Fig. 2C**). As *τ*^2^ controls the between-TR heterogeneity, larger *τ*^2^ would make candidate TRs more easily distinguishable and so lead to better ranking performance (i.e., higher Spearman’s ρ). On the contrary, 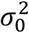 controls the noise level (i.e., how *θ*_*i*,1_ differs from *θ*_*i*′_) for TRs in the second group. Thus, increasing 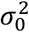 would make the estimation of *θ*_*i*′_ more difficult, reducing the ranking performance.

The number of datasets associated with each TR in the first group was simulated using a lognormal distribution with right-skewness and heavy-tailedness, right truncated at 1,000, which closely mimics the actual count distribution observed in the human reference library (**Fig. 2D**). We also checked the robustness of BIT to the normality assumption in the BIT model. We used two types of non-Gaussian distributions, the Student’s t-distributions (with degrees of freedom 3, 5, and 10) and gamma distributions (shape parameter *α*=3, 5, 10, scale parameter *β*=1), to generate *θ*_*i*_’s and *θ*_*i*′_’s. The overall shapes of these generated distributions are different from Gaussian distributions (**Fig. 2E**). The t-distributions with only a few degrees of freedom are known for heavy-tailedness, and gamma distributions for skewness. The simulated data are then shifted and scaled to recover the same mean and variability for comparison while keeping the heavy-tailedness or skewness. The ranking performance of both methods appears to decline as the deviation from the normality becomes larger. Nevertheless, BIT is much less sensitive and the advantages over the baseline method persist (**Fig. 2F**).

In summary, through simulations in settings mimicking various real scenarios, we showed BIT’s superior performance in accurately estimating parameters and recovering original TR ranks. It consistently outperformed the baseline method, demonstrating high efficiency and robustness even when normality is violated.

### BIT can identify perturbed TRs from differentially accessible regions (DARs)

DARs derived from differential accessibility analysis between two biological conditions are critical in understanding the activity of TRs. Here, we leverage DARs from TR perturbation experiments to validate BIT’s capacity to pinpoint key TRs. The targeted nature of these experiments provides a ground truth, allowing us to assess BIT’s ability to identify the perturbed TRs.

The first experiment is an acute depletion of CTCF in the MLL-rearranged human B cell lymphoblastic leukemia (B-ALL) cell line SEM^40^. The second is a ZBTB7A-knockout experiment in the HUDEP-2 cell line^41^. The third involves a FOXA2-knockout in the pancreatic progenitors 1 (PP1) differentiation stage of human pluripotent stem cells (hPSCs) ^42^. BIT has successfully identified each perturbed TR as one of the top-ranked TRs in all three experiments, and it has also correctly identified several other TRs that contribute to the biological process involved (**Fig. 3A** and see **Tab. S1** for full lists).

**Figure 3.**
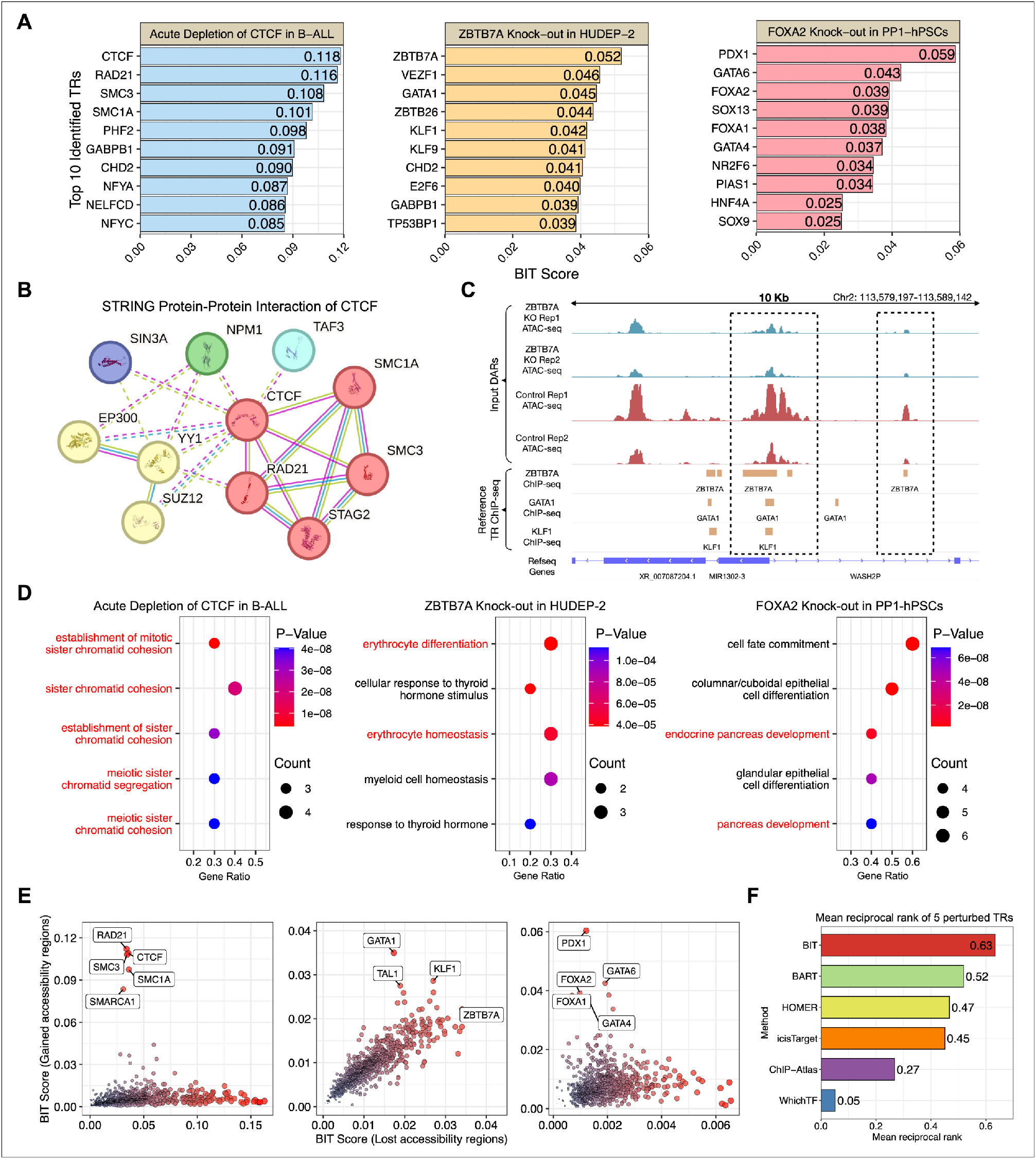
BIT can identify transcriptional regulators (TRs) from differentially accessible regions (DARs). **A**. The top 10 transcriptional regulators (TRs) identified by BIT using DARs from each of the three TR perturbation experiments. **B**. Protein-protein interaction of CTCF derived from STRING. **C**. BIT-identified TRs have binding patterns consistent with the differentially accessible regions observed in TR perturbation experiments. **D**. The top 5 terms ranked by gene ontology enrichment analysis of top TRs in the output list of three cases. **E**. The scatterplot of TRs by using BIT score from gain accessible regions versus loss accessible regions. **F**. Mean reciprocal rank (MRR) of five perturbed TRs for BIT and five state-of-the-art methods (BART, ChIP-Atlas, i-cisTarget, WhichTF, HOMER).

Using DARs from the first experiment as the input regions, the perturbed TR, CTCF, was ranked first. Other top-ranked TRs include RAD21, SMC3, and SMC1A, which are all components of the cohesin complex. SMC proteins and RAD21 are core proteins that form the ring-shaped cohesin complex (cohesin ring) ^43,44^. CTCF and the cohesin ring can be crucial in forming chromatin loops and topologically associating domains (TADs). These two structures are critical in regulating gene transcription^45,46^. Previous studies have confirmed that both CTCF and the cohesin ring are essential for the maintenance of the two structures when using acute depletion assays to handle proteins^47,48^, which can explain why BIT also identified the other three TRs in this study. The result was also validated by the protein-protein interactions of CTCF retrieved from the STRING database^49^, which shows strong associations among CTCF, SMC3, SMC1A, and RAD21 (**Fig. 3B**).

In the second experiment, in addition to the top-ranked ZBTB7A (the perturbed TR), GATA1 and KLF1 were also ranked in the top 10, which have binding patterns consistent with the DARs derived from the experiment (**Fig. 3C**). Early studies have revealed that erythroid differentiation and mutation can be controlled by GATA1-dependent transcription^50,51^, and that ZBTB7A is a cofactor of GATA1 and plays an essential anti-apoptotic role during terminal erythroid differentiation^52^. This explains why GATA1 was also ranked at the top after the knockout of ZBTB7A. In addition, KLF1 has a multifunctional role in erythropoiesis and acts as one of the master regulators^53^.

In the third experiment, FOXA2 ranked third, but notably, other known pancreatic regulators (PDX1, GATA6, FOXA1, and GATA4) were also identified. This is consistent with their established roles in pancreatic development and disease^54-57^. For example, the pioneer factors FOXA1/2 could initiate chromatin accessibility^58^, allowing subsequent binding and transcriptional regulation by PDX1, GATA6, and GATA4.

The results of these three experiments were also validated using Gene Ontology (GO) enrichment analysis, where we used the top TRs identified by BIT to examine enriched biological processes (**Fig. 3D**). The count in this figure indicates the number of TRs associated with each process, and the p-value was calculated using hypergeometric tests to evaluate whether the association of TRs with a biological process is higher than expected by chance. We observed that the top five enriched biological processes, ranked by p-values, include those involving sister chromatid cohesion formation for the first experiment, erythrocyte differentiation and homeostasis for the second experiment, and pancreas differentiation and development for the third experiment. This further supports that BIT was able to identify TRs from DARs.

We further categorized the DARs into two groups based on whether these regions showed increased (gained) or decreased (lost) accessibility after the perturbation of TRs. We separately applied BIT to the two groups and noticed that the perturbed TRs were ranked differently according to their BIT scores (**Fig. 3E**). For example, CTCF and FOXA2 were ranked higher in the group of gained accessibility regions, while ZBTB7A was ranked higher in the group of lost accessibility regions. This provides insights into how the perturbation of these TRs affects chromatin accessibility and demonstrates that BIT can offer additional understanding of the functional role of TRs from the DAR analysis.

We also applied BIT to two more datasets derived from TR knockout in mouse thymocytes to study its performance on data from animal models and validated the results with GO enrichment analysis (**Fig. S3**). BIT not only ranks the perturbed TRs in top positions (no. 3 and no. 2, respectively) but also identifies other experimentally validated TRs strongly associated with perturbed TRs or biological processes involved.

Finally, to compare the performance of BIT with other state-of-the-art methods (BART, ChIP-Atlas, i-cisTarget, WhichTF, and HOMER), we calculated the mean reciprocal rank (MRR) of the five perturbed TRs for each method (**Fig. 3F**). MRR is a commonly used metric to evaluate a method’s ability to rank relevant terms at the top positions^59^. Among the six methods compared, BIT demonstrated the best overall performance.

### BIT can identify cancer-type specific TRs

To further validate BIT’s efficacy, we leveraged cancer-type-specific accessible regions obtained from The Cancer Genome Atlas (TCGA) database^60^. TRs are well-established drivers of crucial cellular processes in various cancers. However, a significant portion of accessible regions recur across multiple cancer types. Including possible non-specific regions can hinder the identification of critical TRs specific to one cancer type. Therefore, employing cancer-type-specific accessible regions offered a more focused approach. We used regions specific to cancers originating from nine tissue types to validate BIT’s performance: breast, bladder, colon, lung, liver, mesothelium, prostate, squamous cell, and testis. More specifically, the ATAC-seq samples used to generate these regions encompassed a broad range of carcinomas, including breast invasive carcinoma (BRCA), bladder urothelial carcinoma (BLCA), and others (see **Supplementary Note 2** for an extended discussion). The identified TRs were cross-validated through existing publications and CRISPR/Cas9 screening data from The Cancer Dependency Map Consortium (DepMap) ^61^, which evaluates cancer cell viability after gene knockout.

First of all, many of BIT’s top-ranked TRs are well-known regulators of normal cell processes such as differentiation and proliferation and have also been implicated in tumor development. A total of 32 TRs (listed by cancer type in **Fig. 4A**) were cross-validated with existing literature (**Tab. S2**), with some already serving as biomarkers or therapeutic targets in cancer treatment^62,63^. For instance, top-ranked TRs in breast cancer included FOXA1, ESR1, and PGR while prostate cancer revealed PIAS1, AR, and HOXB13, and testicular cancer revealed NANOG, SOX17, and POU5F1 (see **Tab. S3** for full lists). Beyond TR identification using estimated BIT scores, BIT also gauges estimation uncertainty. For example, PIAS1 and SOX13 were identified in multiple cancer types, but with higher uncertainty in their BIT score compared to other TRs in the same cancer (**Fig. 4B**). This suggests their potential roles as pan-cancer regulators.

**Figure 4.**
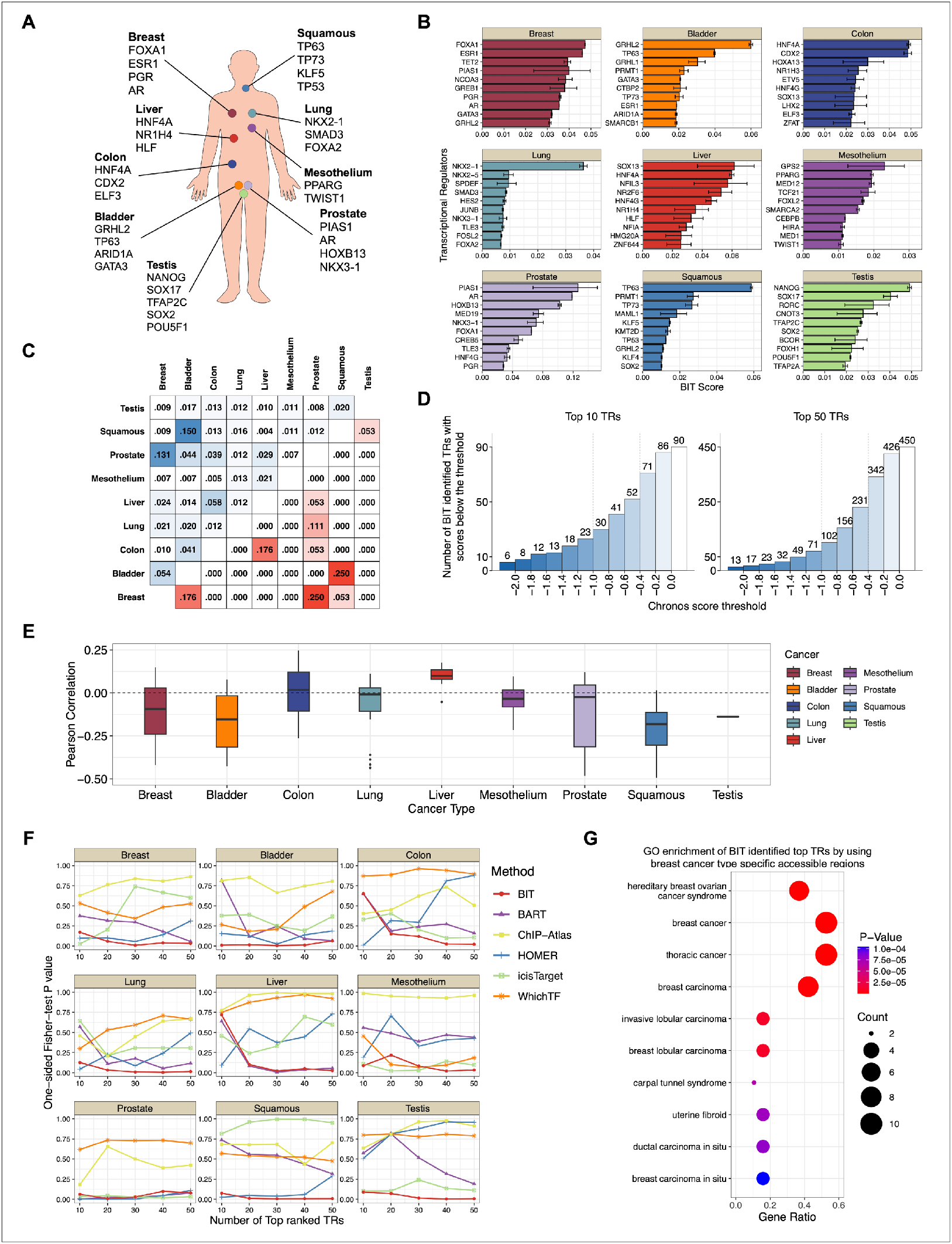
BIT can identify transcriptional regulators from cancer-type specific accessible regions. **A**. 32 TRs identified by BIT from TCGA cancer-type-specific accessible regions validated by existing literature. **B**. BIT scores of top 10 TRs with upper and lower bound of 95% credible intervals. **C**. Jaccard index compares cancer-specific accessible regions between any two cancer types (upper triangle) and compares the sets of top TRs identified by BIT between any two cancer types (lower triangle). **D**. Cumulative number of BIT identified top 10 and top 50 TRs with minimum Chronos scores of ≤−2 to >0. **E**. Boxplots showing the Pearson correlation coefficients between Chronos scores and BIT scores for the top 50 TRs identified by BIT across 209 cell lines from 9 cancer types. **F**. The results of the one-sided Fisher test evaluate whether the top TRs identified by BIT or other methods (BART, ChIP-Atlas, HOMER, i-cisTarget, WhichTF) significantly overlap with functionally essential TRs. **G**. The results of gene-ontology enrichment analysis of top 20 TRs identified by BIT from breast cancer.

Second, we investigated the specificity of BIT-identified TRs across different cancer types. The low Jaccard similarity coefficients (0.003 to 0.150) of the input binary vectors indicated minimal overlap between cancer types (**Fig. 4C**, upper triangle), confirming that retrieved accessible regions are generally specific to their corresponding cancer type. Furthermore, the top-ranked TRs identified by BIT varied significantly across cancer types (**Fig. 4B**), consistent with the generally low Jaccard similarity (**Fig. 4C**, lower triangle). This highlights BIT’s ability to identify key TRs specific to each cancer type.

Third, it is noted that several top-ranked TRs, such as NCOA3 and TET2 in breast cancer, lack known motifs^2^. This absence means these TRs cannot be detected using motif-based methods, despite solid experimental evidence supporting their significant roles in tumor development^64,65^. In contrast, BIT uses TR ChIP-seq data to prioritize these TRs, irrespective of motif information. This highlights BIT’s advantage over traditional motif-based methods, especially when known motifs are ambiguous or absent.

Fourth, given the critical roles of many TRs in cellular viability^7,66,67^, we hypothesized a significant overlap between BIT’s top-ranked TRs and functionally essential TRs (i.e., TRs strongly correlated with cancer cell viability) identified in corresponding cancer types via CRISPR/Cas9 knockout screenings. Indeed, approximately half of the BIT-identified TRs were functionally essential TRs (**Fig. 4D**): 52 out of 90 top-10 TRs and 231 out of 450 top-50 TRs across various cancer types. Here, functionally essential TRs were defined as those with a minimum Chronos (minC) score^61,68^ (across all cell lines) less than -0.4; a more negative minC indicates greater relevance. Furthermore, we computed Pearson correlation coefficients between Chronos and BIT scores of the top 50 TRs identified by BIT in 209 cell lines across the nine cancer types. Boxplots showed in seven out of nine cases, the two scores of most cell lines are negatively correlated, which indicates that higher-ranked TRs generally exhibited lower Chronos scores, thus having a greater impact on cancer cell viability after gene knockout (**Fig. 4E**).

Last but not least, we benchmarked BIT against state-of-the-art methods (e.g., BART, ChIP-Atlas, i-cisTarget, WhichTF, and HOMER) by comparing the enrichment of their top-ranked TRs within functionally essential TRs from CRISPR/Cas9 knockout data. Using a one-sided Fisher exact test, we find that BIT consistently performed at or near the top across all the cancer types with lower p-values (**Fig. 4F**), showing less variation as the number of top TRs increased. While cellular viability is not the sole determinant of TR functionality in cancer, BIT’s robust performance highlights its efficacy in identifying crucial TRs from cancer-type-specific accessible regions, even amidst the multifaceted nature of TR functionality.

In addition to the cross-validation with CRISPR/Cas9 knockout analysis, we conducted several more analyses using various resources: (1) Gene ontology enrichment analysis using BIT’s top TRs identified from breast cancer-specific accessible regions revealed enriched GO terms strongly associated with breast cancer (**Fig. 4G**). (2) We retrieved super-enhancer targeted genes from Cistrome^69^ for six available cancer types, which rank genes based on adjacent super-enhancer activities. We noticed many BIT-identified TRs are also highly ranked by Cistrome (**Fig. S4A**). (3) Kaplan-Meier plotter^70^, a public analysis tool that has collected data from over 30,000 cancer samples, is used to investigate the correlation between gene expression levels and patient survival time. It is observed that multiple BIT-identified TRs show a strong correlation with patient survival time (**Fig. S4B**).

### BIT can identify cell type-specific TRs

The rise of single-cell omics technologies has created a need for computational tools capable of analyzing transcriptional regulation at the individual cell level. scATAC-seq, a powerful technique for profiling chromatin accessibility in single cells, often involves clustering cells and annotating them to specific cell types to overcome the challenge of sparse data. An important task in downstream analyses of scATAC-seq data is to identify key TRs associated with accessible regulatory regions in specific cell types.

In this study, we investigated the application of BIT for identifying crucial TRs using cell type-specific accessible regions derived from scATAC-seq data. Two datasets, including 10K peripheral blood mononuclear cells (PBMCs) and primary liver cancer samples, were used, where the cell types were previously annotated based on the dimension reduction results and marker genes. The UMAP projections of the datasets show a clear separation of different cell types in each dataset (**Fig. 5A**). SnapATAC2^71^ was applied to generate cell type-specific accessible regions. We applied BIT to accessible regions derived from B cells in the PBMCs dataset and malignant cells in the liver cancer dataset. The results were thoroughly validated by different approaches, as described below.

**Figure 5.**
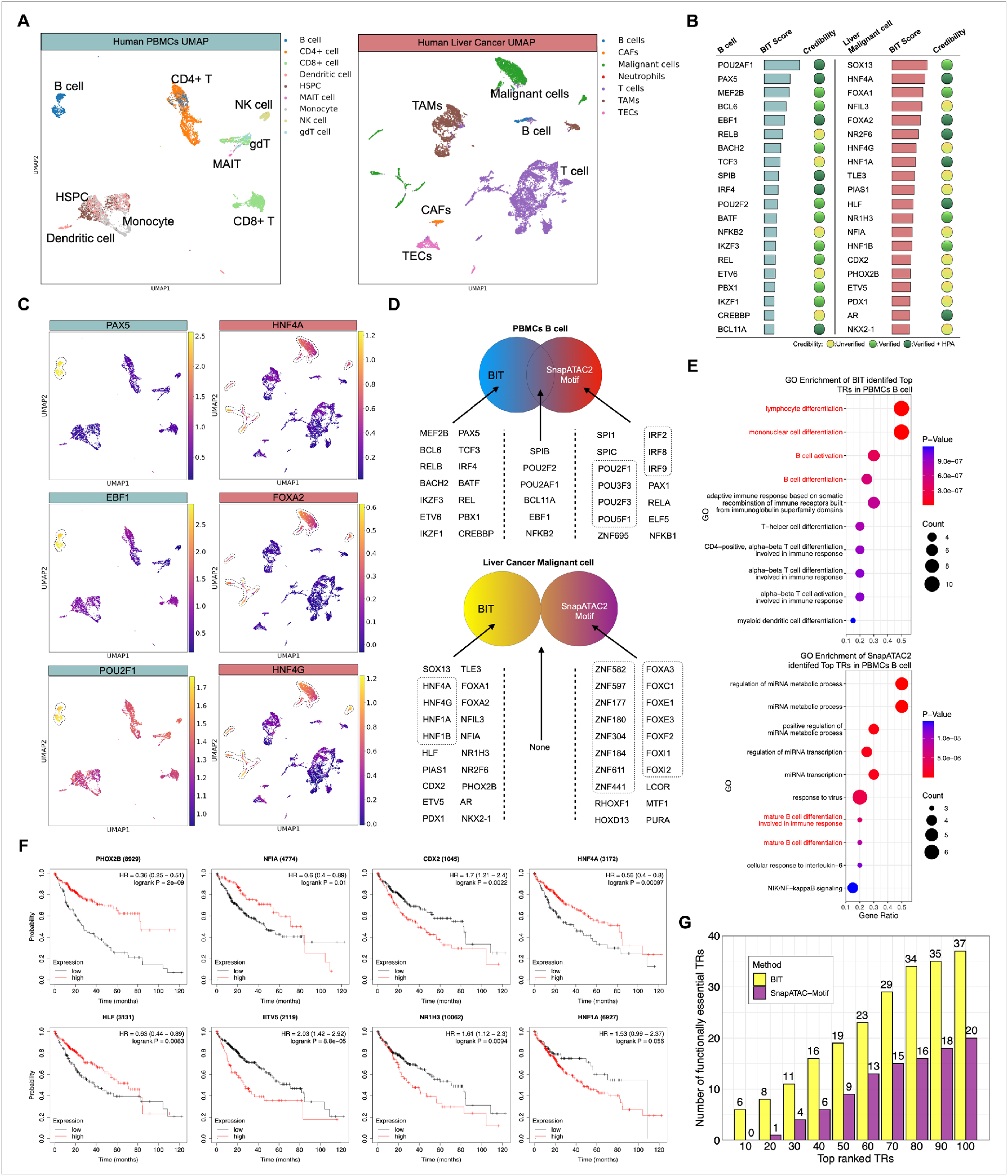
BIT can identify transcriptional regulators from cell-type-specific accessible regions. **A**. Sub-clustering UMAP projection of human Peripheral blood mononuclear cells and primary liver cancer cells. **B**. Top 20 TRs identified by BIT from using B cell-specific and liver malignant cell-specific accessible regions, verified by existing literature and Human Protein Atlas. **C**. UMAP visualization of gene activity of PAX5, EBF1, POU2F1 in PBMCs and FOXA2, HNF4A, and HNF4G in liver cancer samples. **D**. The comparison of top 20 TRs identified by BIT and SnapATAC2. **E**. Gene ontology enrichment results of top 20 TRs identified by BIT and by SnapATAC2 motif enrichment analysis in PBMCs B cells. **F**. Kaplan-Meier Plots of PHOX2B, NFIA, CDX2, HNF4A, HLF, ETV5, NR1H3, HNF1A in liver cancer samples. **G**. Number of BIT and SnapATAC2 identified top 10 to 100 TRs from liver malignant cells with minimum Chronos score ≤−0.4.

First, we evaluated the credibility of the top 20 TRs identified by BIT in the two investigated cell types (**Fig. 5B**) based on existing publications and by examining whether each identified TR has a significantly elevated protein expression in the Human Protein Atlas for the respective cell type (**Tab. S4**). Notably, BIT successfully captured several TRs critical for B cell development. These include PAX5 and EBF1, which were previously reported for their role in maintaining B cell identity. In addition, POU2AF1 and POU2F2 can act as pivotal regulators in B cell proliferation, differentiation, the germinal center transcriptional program, and immune response (see **Tab. S5** for full lists).

For liver cancer malignant cells, we observed consistency between the top TRs identified from scATAC-seq data and those previously found using bulk ATAC-seq data for liver cancer (**Fig. 4B**). Specifically, SOX13, HNF4A, and HLF were again highly ranked by BIT. While the previous analysis used accessible regions from aggregated bulk data, the current analysis leveraged single-cell data specific to malignant cells. This consistency supports the robustness of BIT in identifying critical TRs across different data types.

Second, gene activity plots were generated to demonstrate the high specificity of the identified TRs in both B cells and malignant cells (**Fig. 5C**; left panel for B cells and right panel for malignant cells). Gene activity was assessed by counting the number of Tn5 insertions within the gene’s regulatory regions, a standard approach in scATAC-seq analysis^71,72^. Multiple BIT-identified Top TRs exhibited significantly higher activity in the studied cell types than other cell types, including PAX5, EBF1, and POU2F1 in PBMCs B cells and HNF4A, HNF4G, and FOXA2 in Liver cancer malignant cells.

Third, we also compared the top 20 TRs identified by BIT with those identified through motif enrichment analysis implemented in SnapATAC2, where the two sets have a small or no overlap at all. As mentioned in the introduction, the similarity of binding motifs can hinder the differentiation between TRs from the same protein family. This is evident in the top-ranked TRs identified using SnapATAC2, where results of B cells include the POU, IRF, and PAX families, and results of liver cancer malignant cells exhibit enrichment exclusively for the ZNF and FOX families (**Fig. 5D**). It is crucial to note that not all members within these families hold equal importance. Their inclusion in the top rankings solely reflects the inability of motif enrichment analysis to identify which specific TR within the family directly interacts with the motif site in the accessible chromatin regions. In contrast, BIT’s ability to pinpoint specific TRs, even within families with similar binding motifs, offers a distinct advantage in understanding the precise regulatory mechanisms at play in different cell types.

Next, we performed GO enrichment analysis for the top TRs in B cells identified by BIT and by motif enrichment, separately. The results revealed a strong enrichment for biological processes relevant to B cells or mononuclear cells in the BIT-identified TRs. Conversely, the motif enrichment-identified TRs displayed enrichment for the miRNA metabolic process (**Fig. 5E**).

We further explored the potential of BIT-identified TRs from liver malignant cells as prognostic biomarkers. Kaplan-Meier survival analysis demonstrated a statistically significant association between the expression levels of multiple BIT-identified TRs and patient survival time (**Fig. 5F**), suggesting their promise as prognostic biomarkers in liver cancer. In addition, CRISPR/Cas9 knockout experiments in liver cancer cell lines revealed that BIT identified a greater number of functionally essential TRs compared to motif enrichment analysis in SnapATAC2 (**Fig. 5G**).

In addition to the above results for B cells and liver malignant cells, we generated results for monocytes and dendritic cells in the PBMCs (**Fig. S5**) and for tumor-associated macrophages and T cells in the primary liver cancer samples (**Fig. S6**). Collectively, our findings demonstrate the feasibility of using BIT for the downstream analysis of cell type-specific accessible regions in scATAC-seq data. Beyond feasibility, BIT offers valuable insights into potential regulatory mechanisms. This makes it a compelling alternative to the current dominant motif-based enrichment methods used in scATAC-seq analysis.

## Discussion

We proposed a Bayesian hierarchical approach, called BIT, to identify TRs using user-provided epigenomic regions (peaks), often derived from genome-wide epigenomic profiling data. Using various simulation and application studies, we demonstrated the effectiveness of BIT and its advantages over existing state-of-the-art methods.

### Biological Complexities of TR Regulation

Though BIT demonstrated superior performance consistently in various applications, BIT can suffer from the biological complexities of TR regulation, which can reduce the method’s performance. TR binding is inherently dynamic, changing across cell types and time in response to diverse signals and environmental factors^73^. Chromatin structure and epigenetic modification can further complicate this by affecting the binding affinity and specificity^23,74^. In addition, cooperative binding can also change binding patterns as TRs frequently cooperate to regulate gene expression^75^. A large number of interactions between TRs, DNA, and other cellular components challenge accurate prediction and functional interpretation, while the currently gathered context-specific binding profiles from ChIP-seq experiments only represent a very tiny part of all possible experiments.

### Data quality and sequencing biases

Similar to other NGS-based computational methods, data quality is a critical issue that can affect BIT’s performance. Multiple types of sequencing biases, such as GC content bias or PCR amplification bias, can potentially reduce the reliability of the input epigenomic region set and binding profiles generated by NGS experiments^76^. For instance, the Illumina platform is known for a strong GC bias, wherein epigenomic regions with high or low GC content may be over- or under-represented in the sequencing data^77^. Preferential amplification of certain sequences can also lead to the overrepresentation of specific genomic regions^78^. Therefore, the performance of BIT can depend on the quality of the NGS data used.

### Alternative input from epigenomic profiling data

In the manuscript, we primarily used ATAC-seq datasets that profile chromatin accessibility as input data. However, other techniques such as DNase-seq can also profile the chromatin accessibility and serve as input. In addition, alternative epigenomic profiling techniques are available that also correlate with TR binding activity. For example, H3K27ac ChIP-seq data can mark active promoters and enhancers^79^, which are potential TR binding sites. Therefore, these datasets can also be employed to identify TRs regulating the biological processes of interest.

### Computational efficiency

BIT implements a Gibbs sampling scheme to draw posterior samples for thousands of parameters defined within the model. The number of parameters is determined by the available TRs and associated datasets in the reference library. With the growing availability of ChIP-seq experiments, the number of parameters to infer is expected to increase significantly. In addition, it is common for researchers to generate dozens or even hundreds of epigenomic profiling datasets in a single project. Analyzing these datasets using the Gibbs sampler can become increasingly time-consuming. Therefore, alternative approaches such as variational inference^80,81^ are currently being considered for improving computational efficiency.

In summary, BIT offers valuable insights into transcriptional regulation by providing accurate TR identification through a rigorous Bayesian hierarchical model. This approach leverages the wealth of accumulated TR ChIP-seq data, representing the most accurate *in vivo* context-specific binding patterns. With the increasing popularity of epigenomic profiling techniques, particularly bulk and single-cell ATAC-seq, BIT addresses a critical need for effective tools to analyze this rapidly increasing omics data.

## Methods

### Preprocessing epigenomic NGS data

#### Bulk data

In the TR perturbation-derived DARs analysis, raw ATAC-seq sequence reads were first cleaned using Trim-galore. Read quality was checked by FastQC. Paired-end reads were mapped to the human genome (hg38) or mouse genome (mm10) using bowtie2 applying parameters “bowtie2 --very-sensitive -X 2000”. SAMtools was used for the following steps (i) convert SAM files to BAM files, (ii) remove unmapped reads and duplicates, (iii) generate index files for BAM files. Peaks were called using MACS2. BAM files, index files, and peak files were used for DAR analysis using DiffBind. Reference TR ChIP-seq datasets were retrieved from GTRD, datasets with too low number peaks (<500) and individual peaks with low q-value (<10) were removed. Cancer-type-specific accessible regions were directly retrieved from TCGA.

#### Single-cell data

10K PBMCs and primary liver cancer scATAC-seq fragment files were retrieved from the 10XGenomics and GEO archive. The cell-type-specific accessible regions were generated from each dataset using SnapATAC2. When applicable, the cell type annotations were also retrieved and applied. We also used the SnapATAC2 implemented motif enrichment analysis method to contrast the results.

#### Region mapping

BIT only requires user-provided epigenomic profiling data as input, while leveraging a comprehensive reference library of TR binding profiles from a large collection of TR ChIP-seq datasets. All binding sites (peaks) from a single dataset are mapped to disjoint, consecutive, pre-defined bins of 1000 bps over the entire genome. Each bin is assigned either “0” or “1” according to whether the summit of the peak falls into the bin (in case that the summit information of a peak is missing, the middle point is used instead). Such mapping is also done to the user-provided epigenomic regions. Thus, all data (input and references) are transformed into binary vectors of the same length. We then compare the binary vector from the input with that from each of the TR ChIP-seq datasets, where we define three conditions for any given bin: (i) “matching” if (1,1); (2) “mismatching” if (1,0) or (0,1); and (3) “non-informative” if (0,0). In all input vs. reference comparisons, the first two cases (“informative” bins) are much fewer than the third case (“non-informative” bins) and we drop the third case to mitigate the sparsity while not losing much useful information. Note that the proportion of matching bins among all informative bins is the Jaccard Index, a measure of similarity between two binary vectors.

### The data model of BIT

BIT is the first model-based method that formally distinguishes TRs with multiple datasets (or binding profiles) from those with only a single dataset (or binding profile) in the reference database, to account for both within-TR and between-TR heterogeneity and allow for information borrowing across different TRs and datasets via a hierarchical setup. Our modeling choices are made according to the principle of parsimony in statistics. Thus, BIT strikes a balance between clarity and effectiveness in addressing complex biological questions. It aims to reduce the risk of overfitting while preserving the key insights needed for accurate predictions. An exploratory analysis using the three human TR perturbation datasets reveals the following: (1) When grouped by the same TR, the distribution of the log odds (for matching among informative bins) often exhibits a roughly bell shape, with different groups (TRs) showing different centers and spreads. (2) When calculating the (average) log odds for each TR and categorizing these quantities based on whether the TRs have single or multiple ChIP-seq datasets, the distributions of the two categories exhibit similar centers but different variabilities (**Fig. 6**).

**Figure 6.**
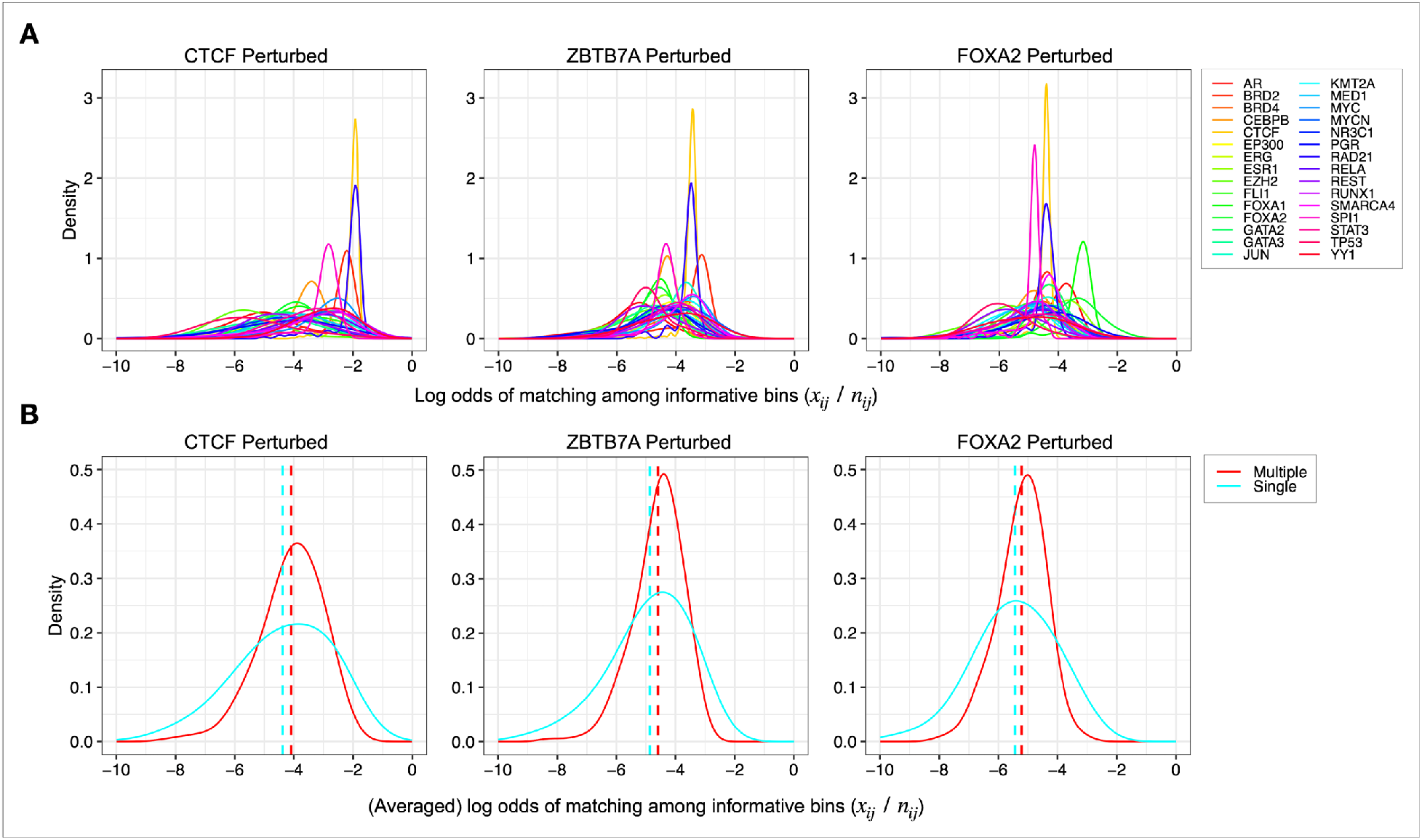
Exploratory analysis using ATAC-seq data from three TR perturbation experiments. **A**. The distributions of log odds of “matching” bins among “informative” bins for the 30 TRs with the highest numbers of TR ChIP-seq datasets available in the reference database. **B**. The distributions of the (average) log odds for all TRs, where TRs are categorized into two groups based on whether each TR has single or multiple TR ChIP-seq datasets.

We index TRs with multiple reference datasets by *i, i* = 1,2, …, ℳ, each with *j* = 1,2, …, *J*_*i*_ datasets. TRs with a single reference dataset are indexed by *i*′ = 1,2, …, ℳ^*c*^, all with *j* ≡ 1 dataset. Let *x*_*ij*_ (*x*_*i*′1_) be the number of “matching” bins out of *n*_*ij*_ (*n*_*i*′1_) “informative” bins by comparing the vectors between the user input and the *j*th reference dataset of the *i*th TR (or the only dataset of the *i* ′ th TR). The probability of a bin being “matching” given it is “informative” is denoted by *p*_*ij*_ (*p*_*i*,1_) such that *x*_*ij*_ ∼ *Binom*(*n*_*ij*_, *p*_*ij*_) and *x*_*i*′1_ ∼ *Binom*(*n*_*i*′1_, *p*_*i*′1_). Let *θ*_*ij*_ ≡ *logit*(*p*_*ij*_) (*θ*_*i*′1_ ≡ *logit*(*p*_*i*′1_)) be the log odds of matching among informative bins, which can be interpreted as the Jaccard similarity index (or simply, the importance score) at the dataset level. We further model 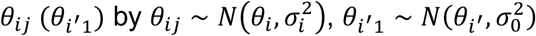 and all *θ*_*i*_, *θ*_*i*′_ ∼ *N*(*μ, τ*^2^), where *μ* and *τ*^2^ represent the global mean and variability of TR-level importance scores. In this way, for each TR with multiple datasets, we model its importance scores by a normal distribution with distinctive mean *θ*_*i*_ and variance 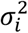, reflecting its overall importance and within-TR heterogeneity, respectively (**Fig. 6A**); all TRs with only one dataset share a common variance 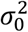 for information pooling; and all TRs, no matter which group they are in, share a common mean *μ* (**Fig. 6B**).

It turns out that the entire hierarchical model (**Fig. 1B**) is simple yet effective, as shown in our simulation and applications:

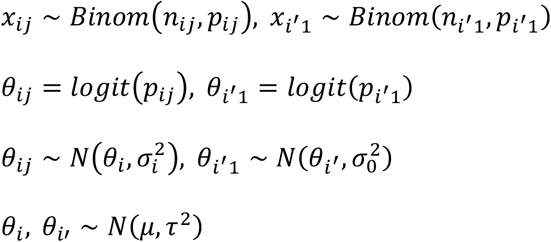

Unlike many existing algorithms^31,32,39^, BIT inherently integrates information from reference datasets of different TRs to assign one final rank of each TR based on the TR-level importance score *θ*_*i*_ (*θ*_*i*′_) rather than conducting thousands of separate *ad-hoc* statistical tests.

The (hyper)parameters of our BIT model are collectively denoted by 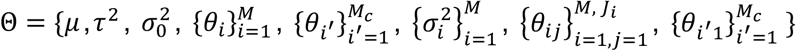 and observed data are 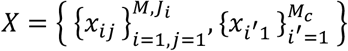.

Then the joint distribution of (X, Θ) can be written as

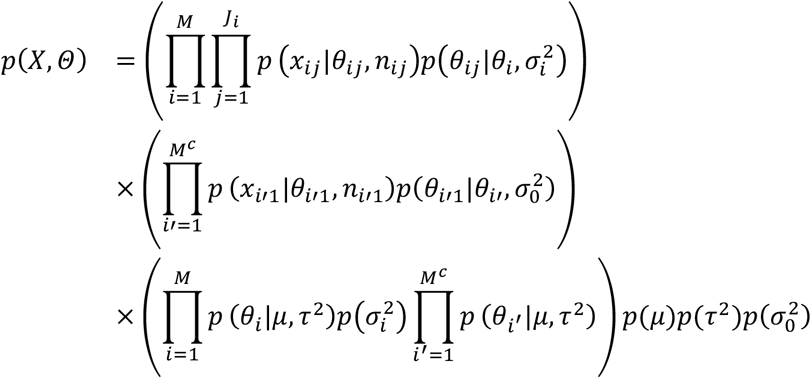

where

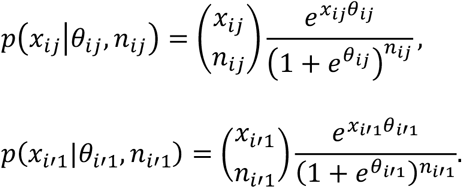

### Prior specification

The global location parameter *μ* and all variance parameters including 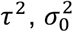 and 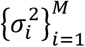 are assumed *a priori* independent. For *μ*, we consider the non-informative flat prior *μ* ∼ *uniform*(*L* _μ_, *U*_μ_), which should provide a sufficiently wide coverage for all plausible values of *μ* suggested by data. Detailed discussion about how to set up the upper and lower bounds can be found in Jia et al. ^82^ For each variance parameter involved, we specify an inverse gamma prior distribution *IG*(*a, b*), where *a* and *b* are tiny positive values to make prior very vague and diffuse (e.g., *a* = *b* = 0.01).

### Pólya-gamma data augmentation and Gibbs sampler

BIT uses a Gibbs sampler, an efficient Monte Carlo Markov chain (MCMC) algorithm to generate posterior samples from the joint probability distribution *p*(*Θ*|*x*) that is proportional to *p*(*x, Θ*). We proceed to derive the full conditionals to set up the Gibbs sampler efficiently. The full conditional posterior distribution of each parameter can be written as *p*(*θ*|*x, Θ* \ *θ*), where *Θ* \ *θ* denote the set of parameters after removing θ from *Θ*. For 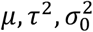,and 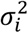, the conditional posterior distributions are known distributions which can be directly sampled, given below:

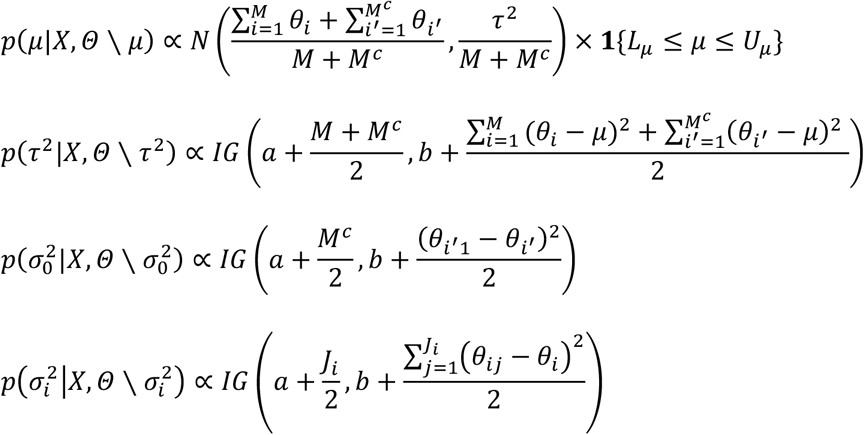

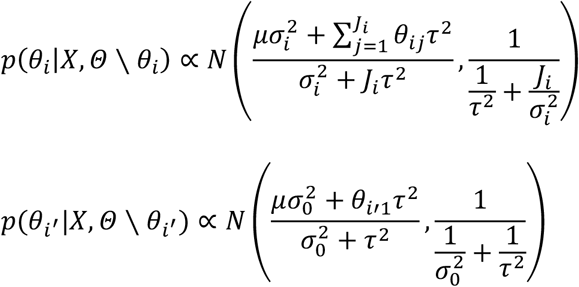

However, it is difficult to sample from the conditional posteriors of *p*(*θ*_*ij*_|*x, Θ* \ *θ*_*ij*_) and *p*(*θ*_*i*,1_|*x, Θ* \ *θ*_*i*,1_) as these are intractable. We employ a data augmentation strategy based on Pólya-Gamma (PG) latent variables^83^. Specifically, we define auxiliary variables λ_*ij*_ and λ_*i* ′1_, which follow the Pólya-Gamma distribution.

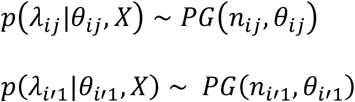

Then the conditionals given the defined auxiliary parameters can be easily written as the form of known distribution.

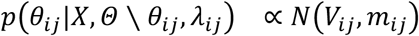

where

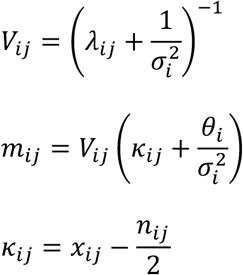

Similarly, we can derive the conditional posterior of *θ*_*i* ′1_. Given that all conditionals are now known distributions, we can design an efficient Gibbs sampler in which all quantities are drawn sequentially and generated readily without using any built-in sampling algorithms (such as Metropolis-Hastings or Acceptance/Rejection sampling) that can greatly slow down the computation.

### Bayesian inference and TR rankings

Inference about TR-level importance parameters can be made by marginalizing over posterior samples generated from the Gibbs sampler. Let 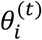 and 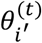 be the posterior draw of *θ*_*i*_ and *θ*_*i* ′_in the *t* th iteration of MCMC after the burn-in period, where *t* = 1, …, *T*. Then *θ*_*i*_ and *θ*_*i*′_ are estimated by 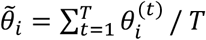 and 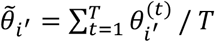. Then, these logit-transformed probabilities 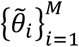 and 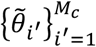 can be used to produce a final ranked TR list or to be further transformed back to the 0-1 scale. To gauge prediction uncertainty, credible intervals of any parameter of interest can be easily obtained from percentiles of draws or as highest posterior density intervals.

### The “naïve” method as a baseline for comparison in simulation

To compare the performance of BIT in estimating model parameters, we used a naïve method as the baseline, where *μ* is estimate by

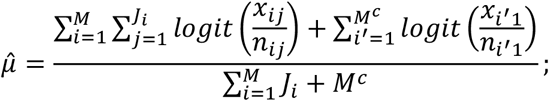

*τ*^2^is estimated by

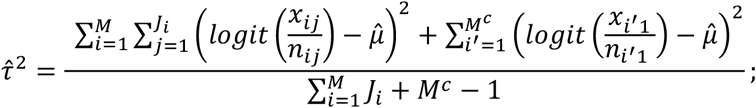

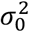 is estimated by

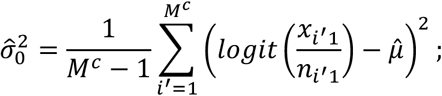

and *θ*_*i*_ and *θ*_*i* ′_ are estimated by

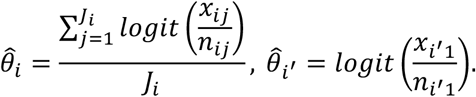

In the unlikely case that 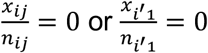, we add a tiny positive value 0.0001 to the zeros.

### BIT validation analyses

#### Computation of mean reciprocal rank

Let the rank of the perturbed TR in the *i*^*th*^ output list generated by a method be denoted as *R*_*i*_, the mean reciprocal rank (MRR) of the five perturbed TRs (CTCF, ZBTB7A, FOXA2, RUNX1, BCL11B) was computed as:

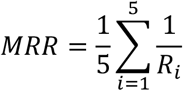

We computed the metric separately for the six methods (BIT, BART, ChIP-Atlas, i-cisTarget, WhichTF, and HOMER).

#### Cross-validation with CRISPR/Cas9 screening data

Data of TR knockout effects on viability of cell lines were retrieved from DepMap. We filtered out TRs that are not included in the BIT database or the DepMap data. Cell lines under the same lineage are grouped to find TRs with a Chronos score lower than -0.4, which are classified as functionally essential TRs. 2-by-2 contingency tables were created based on whether top-ranked TRs identified by each method are also functionally essential TRs. Finally, one-sided Fisher exact tests were applied to each table to compare the capability of methods to identify TRs that are also functionally essential TRs in each cancer type.

To compare with other state-of-the-art methods, the original implementation of BART, HOMER, and WhichTF were retrieved from https://github.com/zanglab/bart2, http://homer.ucsd.edu/homer/, and https://bitbucket.org/bejerano/whichtf. For ChIP-Atlas and i-cisTarget, we used online portals. The same epigenomic profiling datasets given to BIT were used as input, and output TR ranking lists were used to perform Fisher-exact tests with the same steps as for BIT.

#### Gene ontology enrichment analysis

We used the R package “clusterProfiler” to conduct gene ontology enrichment analysis. The top-ranked TRs in the output list of BIT were first converted to their ensemble ids, next by contrasting them to the universe of all available genes. We computed the significance of the enrichment of the top TRs in the GO terms. The top 10 TRs in TR perturbation experiments, and top 20 TRs in the remaining cases were used. We used q-value cutoff of 0.05 and listed the top 5 or 10 terms in the final outputs.

## Data Availability

Only public datasets were used in this study. The reference ChIP-seq datasets were retrieved from GTRD. TR perturbation experiment data were retrieved from GEO (GSE153237^40^, GSE173416^41^, GSE114102^42^, and GSE234331^84^). Cancer-type specific accessible regions were retrieved from TCGA^60^. CRISPR/Cas9 knockout screening data were retrieved from DepMap^61^ (https://depmap.org/portal/). scATAC-seq data were retrieved from 10XGenomics and GSE227265^85^. We initialized a data repository on Zenodo (https://zenodo.org/records/13732877).

## Code Availability

BIT software is available on GitHub (https://github.com/ZeyuL01/BIT), and through a web portal for online analysis (http://43.135.174.109:8080/).

## Acknowledgements

This work was supported by the following funding: the Rally Foundation, Sam Day Foundation, Children’s Cancer Fund (Dallas), the Cancer Prevention and Research Institute of Texas (RP180319, RP200103, RP220032, RP170152 and RP180805), and the National Institutes of Health funds (R21CA259771, P30CA142543, R01HG011996, and R01HL144969) (to L.X.) (R01CA258584) (to X.W.)

## Contributions

Lu, Z. conducted data-processing, model design, simulation, and validation.

Xu, L. and Wang, X. conceived the ideas, designed the study, and acquired the funding.

All members have read, revised, and approved the final manuscript.

## Corresponding authors

Correspondence to Xinlei Wang or Lin Xu.

