## Supplementary material for "BIT: Bayesian Identification of Transcriptional Regulators from Epigenomics-Based Query Region Sets": Fig. S1

This file includes **supplementary figures S1-S6**, **supplementary notes 1-2**.

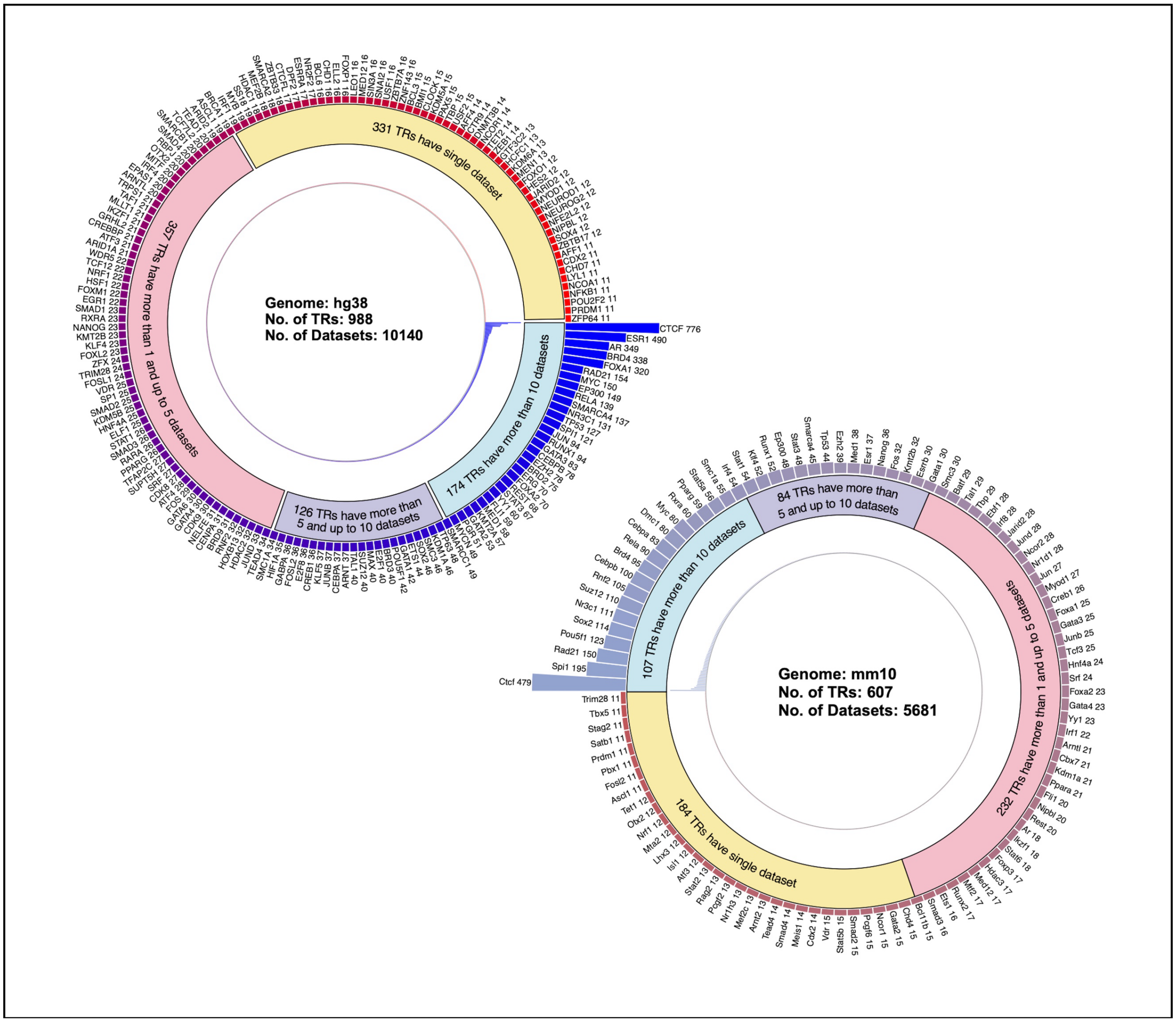


**Figure S1. The BIT pre-processed TR ChIP-seq profiles in the reference library for genome hg38 and mm10.** The outer layer plots the TRs with more than 10 TR ChIP-seq datasets available, the middle layer shows the partition of TRs categorized into four classes based on number of datasets, and the inner layer displays all TRs associated datasets.

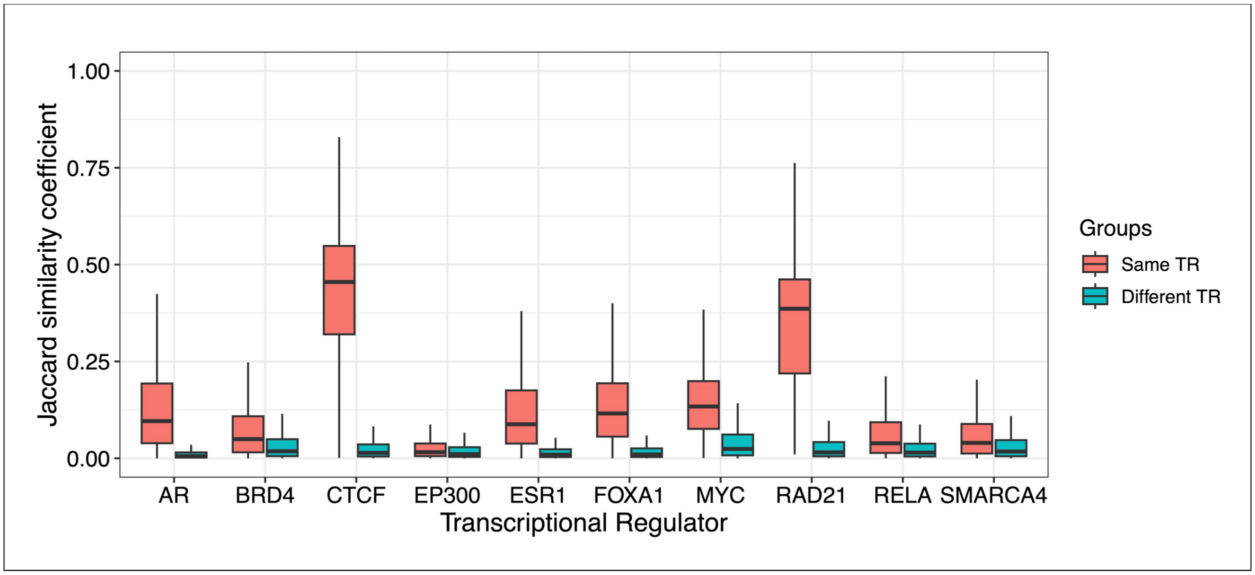


**Figure S2. Boxplots of Jaccard similarity coefficients between two TR binding profiles from ChIP-seq datasets of the same TR (red) vs. different TR's (green).** For example, for the TR named AR, the comparison involves calculation of Jaccard similarity coefficients for binary vectors derived from two AR ChIP-seq datasets vs. binary vectors derived from an AR ChIP-seq dataset and the other randomly sampled from non-AR ChIP-seq datasets. Results for the 10 TRs with the largest number of TR ChIP-seq dataset are reported. Clearly, binding patterns of the same TR across different conditions be more similar compared to those of different TRs.

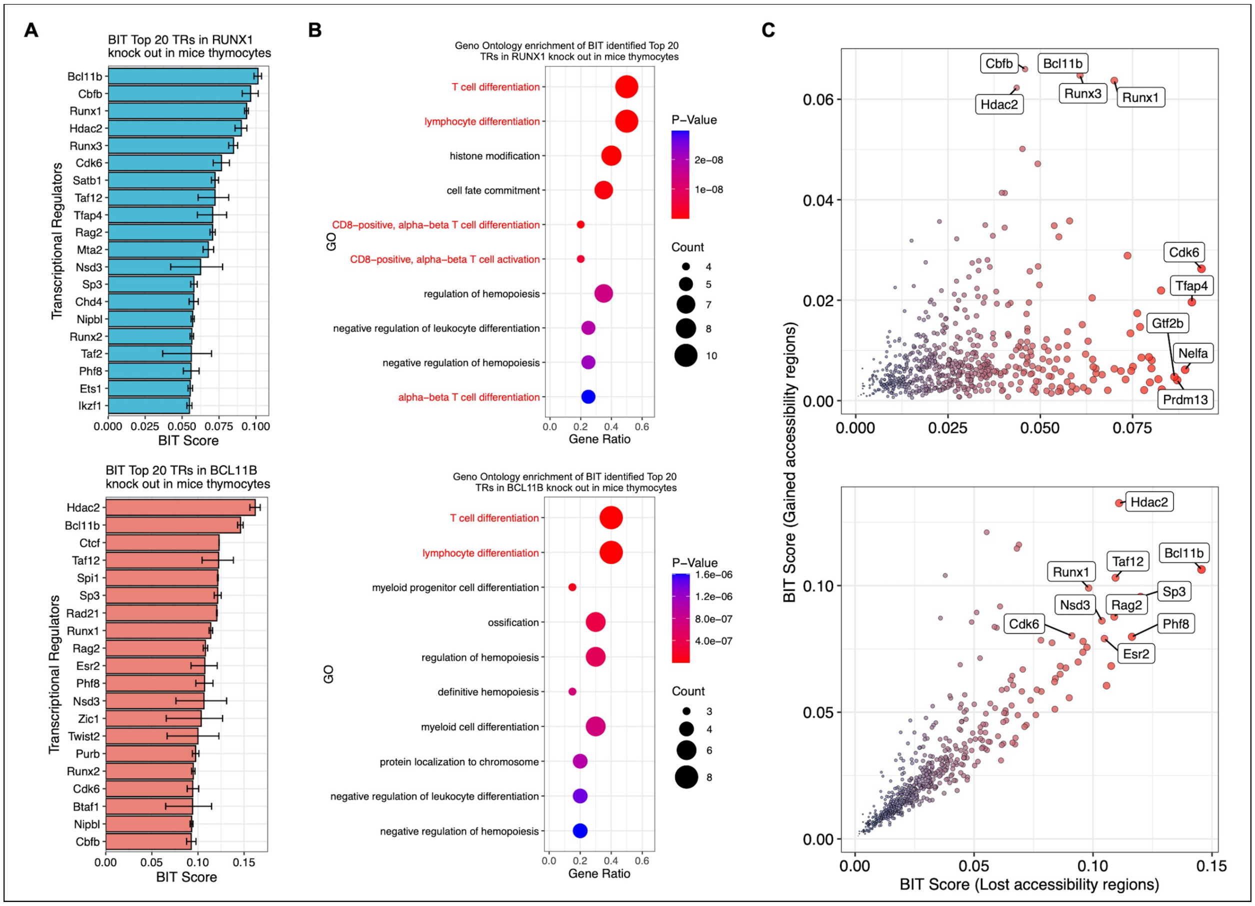


**Figure S3. BIT validation using differentially accessible regions generated from mouse models (mm10). A.** BIT-identified Top 20 TRs from RUNX1 and BCL11B knock out experiments conducted in mice thymocytes. **B.** Gene ontology enrichment results of the Top 20 TRs in the two knock out cases. **C.** BIT Scores of TRs identified from gained accessibility regions vs. those from lost accessibility regions after the knockout of RUNX1 and BCL11B, respectively.

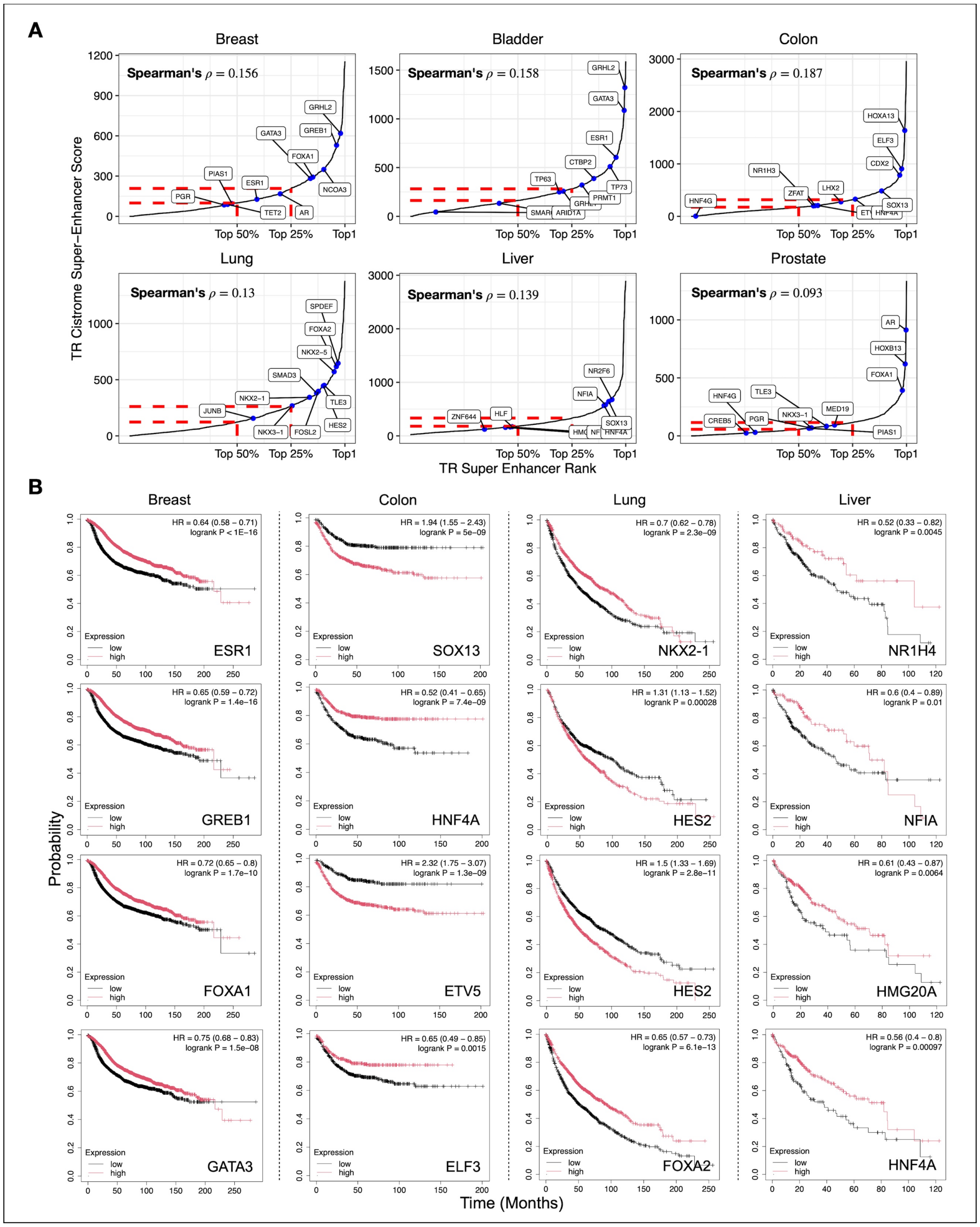


**Figure S4. Extended validation of BIT identified transcriptional regulators (TRs) from cancer-type-specific accessible regions. A.** The Cistrome super-enhancer scores of BIT identified top TRs in six available cancer types. The locations of top 10 BIT identified TRs are marked with blue dots, showing most of them are among the top 25% and nearly all are among the top 50%. Spearman's $\rho$ values between the TR ranks in the BIT and Cistrome super-enhancer lists are consistently positive for all six cancers. **B.** Kaplan-Meier plots of 16 BIT identified top TRs in 4 different cancer types. P-value was calculated using the Mantel-Cox test.

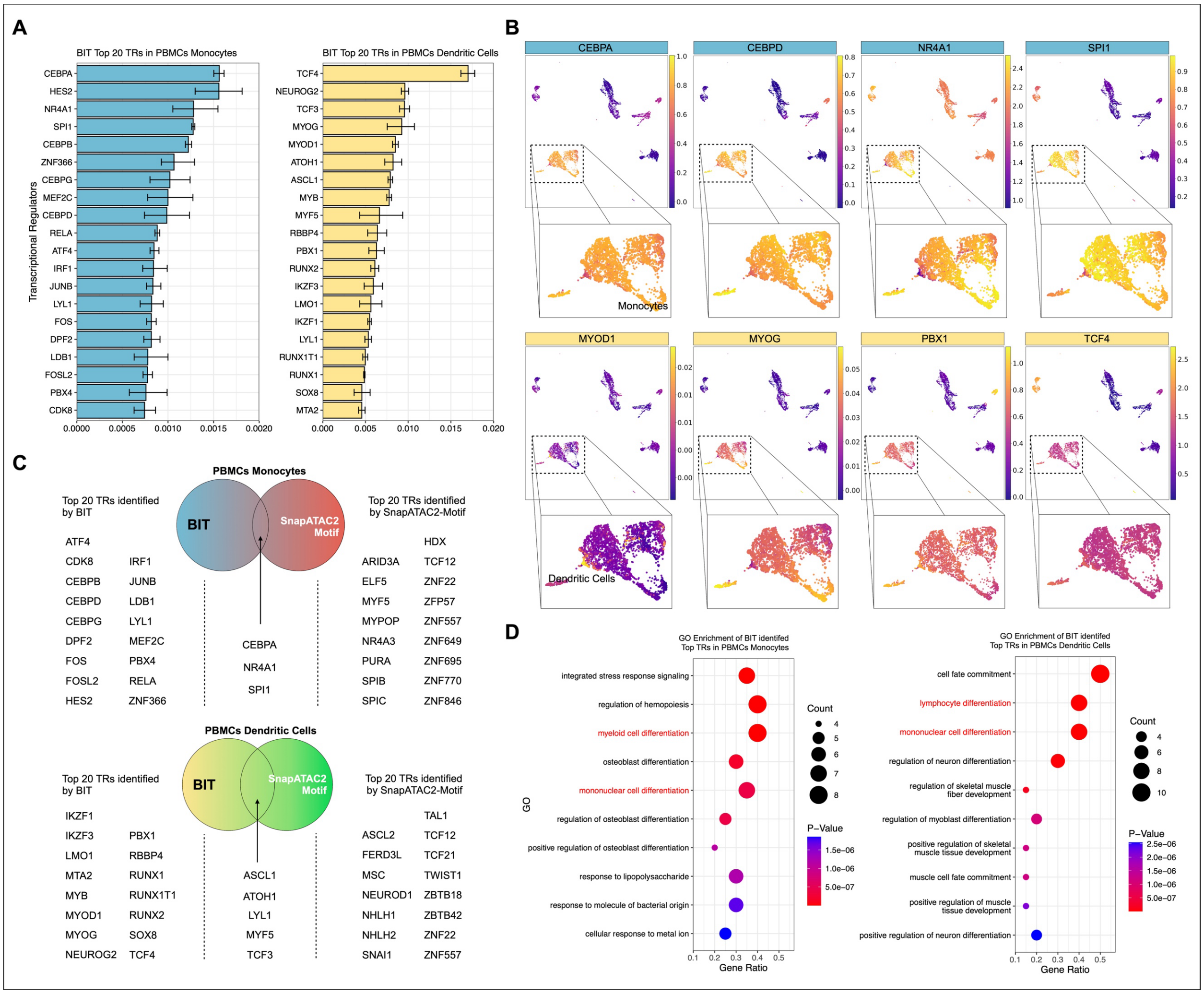


**Figure S5. BIT can identify critical TRs in monocytes and dendritic cells from PBMCs. A.** Top 20 BIT identified TRs in PBMCs monocytes and dendritic cells. **B.** Gene activity of CEBPA, CEBPD, NR4A1, SPI1, MYOD1, MYOG, PBX1, and TCF4 in PBMCs. **C.** Comparison of Top 20 BIT identified TRs versus Top 20 TRs identified by SnapATAC2 using motif enrichment methods. **D.** The GO enrichment results of BIT identified top TRs.

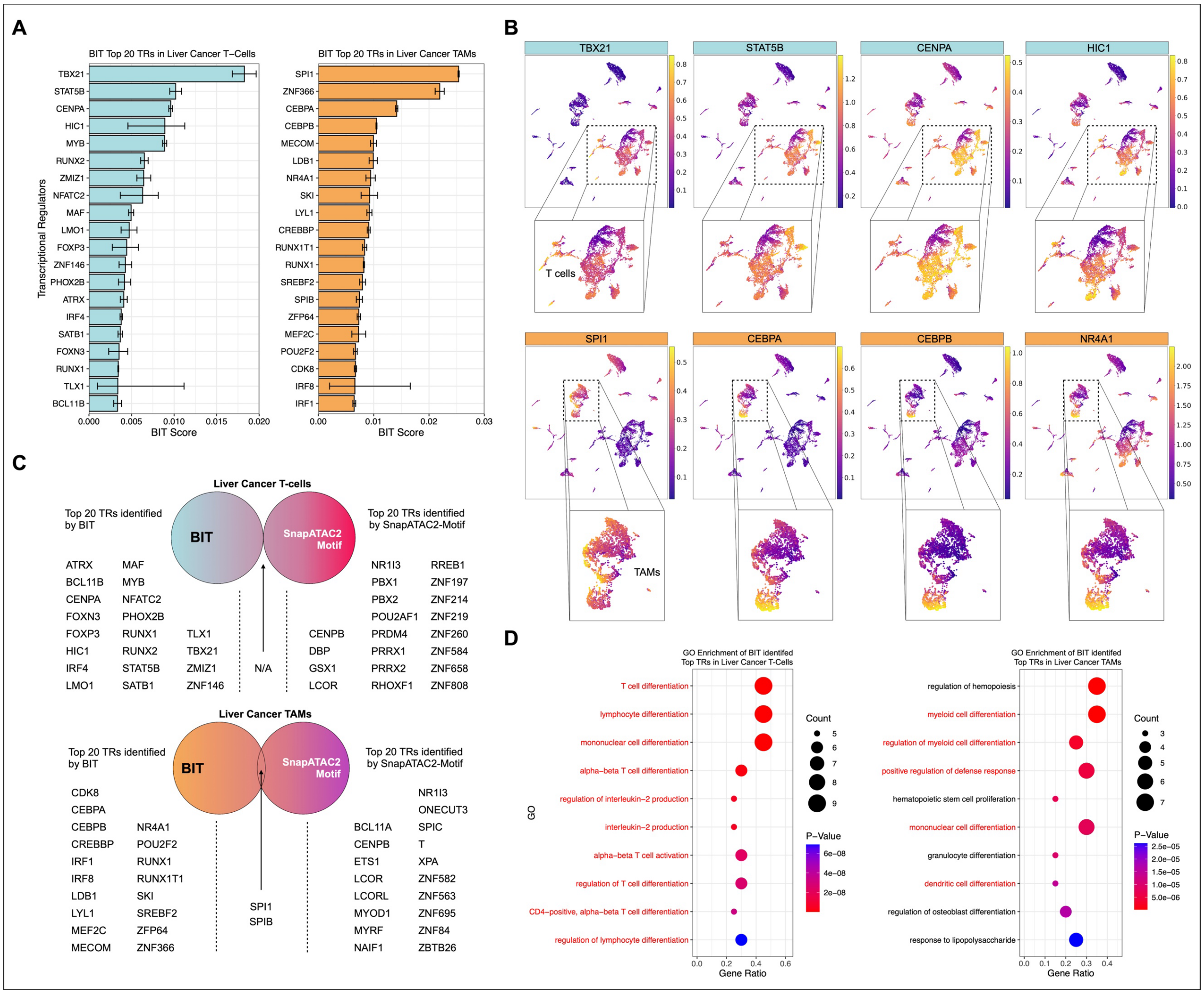


**Figure S6. BIT can identify critical TRs in liver cancer T-cells and Tumor-Associated Macrophages (TAMs). A.** Top 20 BIT identified TRs in liver cancer T-cells and TAMs. **B.** Gene activity of TBX21, STAT5B, CENPA, HIC1, SPI1, CEBPA, CEBPB, and NR4A1 in Liver cancer cells. **C.** Comparison of Top 20 BIT identified TRs versus Top 20 TRs identified by SnapATAC2 using motif enrichment methods. **D.** The GO enrichment results of BIT identified top TRs.

### Supplementary Note 1.

#### S1.1. Notation

| Notation | | | |
|---|---|---|---|
| $i$ | Index of TR with multiple reference datasets. | $\theta_i(\theta_{i'})$ | The TR level importance score. |
| $i'$ | Index of TR with a single reference dataset. | $\sigma_i^2$ | Variance across different datasets of the $i^{th}$ TR. |
| $j$ | Index of reference dataset. | $\sigma_0^2$ | Variance for TRs with only one dataset. |
| $J_i$ | Total number of reference datasets of the $i^{th}$ TR. | $\mu$ | Global mean of TR-level importance. |
| $\mathcal{M}$ | Total number of TRs with multiple datasets. | $\tau^2$ | Global variability of TR-level importance. |
| $\mathcal{M}^c$ | Total number of TRs with a single dataset. | $\lambda_{ij}(\lambda_{i'1})$ | Auxiliary variables follow Pólya-gamma distributions. |
| $x_{ij}(x_{i'1})$ | Number of "matching" bins. | $\kappa_{ij}(\kappa_{i'1})$ | Auxiliary constant for the data augmentation. |
| $n_{ij}(n_{i'1})$ | Number of "informative" bins. | $L_\mu, U_\mu$ | Lower and upper bound of uniform prior of $\mu$. |
| $p_{ij}(p_{i'1})$ | The probability of a "bin" being matching given it is informative. | $a, b$ | Tiny positive values for inverse-gamma prior of $\tau^2$, $\sigma_0^2$, and $\sigma_i^2$. |
| $\theta_{ij}(\theta_{i'1})$ | Log odds of matching among informative bins. | | |

#### S1.2. Derivation of full conditionals

The (hyper) parameters of our BIT model are collectively denoted by $\Theta = \{\mu, \tau^2, \sigma_0^2, \{\theta_i\}_{i=1}^{M},$ $\{\theta_{i'}\}_{i'=1}^{M_c}, \{\sigma_i^2\}_{i=1}^{M}, \{\theta_{ij}\}_{i=1,j=1}^{M,J_i}, \{\theta_{i'1}\}_{i'=1}^{M_c}\}$ and observed data are $X = \left\{ \{x_{ij}\}_{i=1,j=1}^{M,J_i}, \{x_{i'1}\}_{i'=1}^{M_c} \right\}$.

Then the joint distribution of $(X, \Theta)$ can be written as

$$
\begin{aligned}
p(X,\Theta) \quad &= \left(\prod_{i=1}^{M}\prod_{j=1}^{J_i} p\left(x_{ij}|\theta_{ij},n_{ij}\right)p\left(\theta_{ij}|\theta_i,\sigma_i^2\right)\right)\\
&\times\left(\prod_{i\prime=1}^{M^c} p\left(x_{i\prime 1}|\theta_{i\prime 1},n_{i\prime 1}\right)p(\theta_{i\prime 1}|\theta_{i\prime},\sigma_0^2)\right)\\
&\times\left(\prod_{i=1}^{M} p\left(\theta_i|\mu,\tau^2\right)p(\sigma_i^2)\prod_{i\prime=1}^{M^c} p\left(\theta_{i\prime}|\mu,\tau^2\right)\right)p(\mu)p(\tau^2)\text{[illegible]}(\sigma_0^2)
\end{aligned}
$$

so that

$$
\begin{aligned}
p(\Theta|X) \quad &\propto \left(\prod_{i=1}^{M}\prod_{j=1}^{J_i}\binom{n_{ij}}{x_{ij}}\frac{\left(e^{\theta_{ij}}\right)^{x_{ij}}}{\left(1+e^{\theta_{ij}}\right)^{n_{ij}}}\times\left(\sigma_i^2\right)^{-\frac{1}{2}}\exp\left(-\frac{\left(\theta_{ij}-\theta_i\right)^2}{2\sigma_i^2}\right)\right)\\
&\times\left(\prod_{i'=1}^{M^c}\binom{n_{i'1}}{x_{i\prime 1}}\frac{\left(e^{\theta_{i'1}}\right)^{x_{i'1}}}{\left(1+e^{\theta_{i'1}}\right)^{n_{i'1}}}\times(\sigma_0^2)^{-\frac{1}{2}}\exp\left(-\frac{(\theta_{i'1}-\theta_{i'})^2}{2\sigma_0^2}\right)\right)\\
&\times\left(\prod_{i=1}^{M}(\tau^2)^{-\frac{1}{2}}\exp\left(-\frac{(\theta_i-\mu)^2}{2\tau^2}\right)\left(\sigma_i^2\right)^{-a}\exp\left(-\frac{b}{\sigma_i^2}\right)\prod_{i'=1}^{M^c}(\tau^2)^{-\frac{1}{2}}\exp\left(-\frac{(\theta_{i'}-\mu)^2}{2\tau^2}\right)\right)\\
&\times\mathbf{1}\left\{L_\mu\le\mu\le U_\mu\right\}\times(\tau^2)^{-a}\exp\left(-\frac{b}{\tau^2}\right)\times(\sigma_0^2)^{-a}\exp\left(-\frac{b}{\sigma_0^2}\right)
\end{aligned}
$$

Therefore, the conditional posterior of each parameter can be derived as:

(1) $p(\mu|X,\Theta\setminus\mu)$

$$
\begin{aligned}
p(\mu|X,\Theta\setminus\mu) &\propto \prod_{i=1}^{M}(\tau^2)^{-\frac{1}{2}}\exp\left(-\frac{(\theta_i-\mu)^2}{2\tau^2}\right)\prod_{i'=1}^{M^c}(\tau^2)^{-\frac{1}{2}}\exp\left(-\frac{\left(\theta_{i'}-\mu\right)^2}{2\tau^2}\right)\\
&\propto \exp\left(-\frac{M(\theta_i-\mu)^2+M^c(\theta_{i'}-\mu)^2}{2\tau^2}\right)\times\mathbf{1}\left\{L_\mu\le\mu\le U_\mu\right\}\\
&\propto N\left(\frac{\sum_{i=1}^{M}\theta_i+\sum_{i'=1}^{M^c}\theta_{i'}}{M+M^c},\frac{\tau^2}{M+M^c}\right)\times\mathbf{1}\{L_\mu\le\mu\le U_\mu\}
\end{aligned}
$$

(2) $p(\tau^2|X,\Theta\setminus\tau^2)$

$$p(\tau^2|X,\Theta \setminus \tau^2) \propto \prod_{i=1}^{M} (\tau^2)^{-\frac{1}{2}} \exp\left(-\frac{(\theta_i-\mu)^2}{2\tau^2}\right) \prod_{i'=1}^{M^c} (\tau^2)^{-\frac{1}{2}} \exp\left(-\frac{\left(\theta_{i'}-\mu\right)^2}{2\tau^2}\right) \times (\tau^2)^{-a} \exp\left(-\frac{b}{\tau^2}\right)$$

$$\propto (\tau^2)^{-\left(a+\frac{M+M^c}{2}\right)} \exp\left(-\frac{b+\frac{1}{2}\sum_{i=1}^{M}(\theta_i-\mu)^2+\frac{1}{2}\sum_{i'=1}^{M^c}(\theta_{i'}-\mu)^2}{\tau^2}\right)$$

$$\propto IG\left(a+\frac{M+M^c}{2}, b+\frac{\sum_{i=1}^{M}(\theta_i-\mu)^2+\sum_{i'=1}^{M^c}(\theta_{i'}-\mu)^2}{2}\right)$$

(3) $p(\sigma_0^2|X,\Theta \setminus \sigma_0^2)$

$$p(\sigma_0^2|X,\Theta \setminus \sigma_0^2) \propto (\sigma_0^2)^{-a} \exp\left(-\frac{b}{\sigma_0^2}\right) \prod_{i'=1}^{M^c} (\sigma_0^2)^{-\frac{1}{2}} \exp\left(-\frac{\left(\theta_{i'1}-\theta_{i'}\right)^2}{2\sigma_0^2}\right)$$

$$\propto (\sigma_0^2)^{-\left(a+\frac{M^c}{2}\right)} \exp\left(-\frac{b+\frac{1}{2}\sum_{i'=1}^{M^c}(\theta_{i'1}-\theta_{i'})^2}{\sigma_0^2}\right)$$

$$\propto IG\left(a+\frac{M^c}{2}, b+\frac{\sum_{i'=1}^{M^c}(\theta_{i'1}-\theta_{i'})^2}{2}\right)$$

(4) $p\left(\sigma_i^2\middle|X,\Theta \setminus \sigma_i^2\right)$

$$p\left(\sigma_i^2\middle|X,\Theta \setminus \sigma_i^2\right) \propto (\sigma_i^2)^{-a} \exp\left(-\frac{b}{\sigma_i^2}\right) \prod_{j=1}^{J_i} (\sigma_i^2)^{-\frac{1}{2}} \exp\left(-\frac{\left(\theta_{ij}-\theta_i\right)^2}{2\sigma_i^2}\right)$$

$$\propto \left(\sigma_i^2\right)^{-\left(a+\frac{J_i}{2}\right)} \exp\left(-\frac{b+\frac{1}{2}\sum_{j=1}^{J_i}\left(\theta_{ij}-\theta_i\right)^2}{\sigma_i^2}\right)$$

$$\propto IG\left(a+\frac{J_i}{2}, b+\frac{\sum_{j=1}^{J_i}\left(\theta_{ij}-\theta_i\right)^2}{2}\right)$$

(5) $p(\theta_i|X,\Theta \setminus \theta_i)$

$$p(\theta_i|X,\Theta \setminus \theta_i) \propto (\tau^2)^{-\frac{1}{2}} \exp\left(-\frac{(\theta_i-\mu)^2}{2\tau^2}\right) \times \prod_{j=1}^{J_i} (\sigma_i^2)^{-\frac{1}{2}} \exp\left(-\frac{\left(\theta_{ij}-\theta_i\right)^2}{2\sigma_i^2}\right)$$

$$\propto \exp\left(-\frac{(\theta_i-\mu)^2}{2\tau^2}-\frac{\sum_{j=1}^{J_i}\left(\theta_{ij}-\theta_i\right)^2}{2\sigma_i^2}\right)$$

$$\propto \exp\left(-\frac{\sigma_i^2(\theta_i-\mu)^2+\tau^2\sum_{j=1}^{J_i}\left(\theta_{ij}-\theta_i\right)^2}{2\tau^2\sigma_i^2}\right)$$

$$\propto \exp\left(-\frac{\left(\theta_i-\frac{\mu\sigma_i^2+\sum_{j=1}^{J_i}\theta_{ij}\tau^2}{\sigma_i^2+J_i\tau^2}\right)^2}{\frac{2\tau^2\sigma_i^2}{\sigma_i^2+J_i\tau^2}}\right)$$

$$\propto N\left(\frac{\mu\sigma_i^2+\sum_{j=1}^{J_i}\theta_{ij}\tau^2}{\sigma_i^2+J_i\tau^2},\frac{1}{\frac{1}{\tau^2}+\frac{J_i}{\sigma_i^2}}\right)$$

(6) $p(\theta_{i'}|X,\Theta\setminus\theta_{i'})$

$$p(\theta_{i'}|X,\Theta\setminus\theta_{i'})\propto(\tau^2)^{-\frac{1}{2}}\exp\left(-\frac{\left(\theta_{i'}-\mu\right)^2}{2\tau^2}\right)\times(\sigma_0^2)^{-\frac{1}{2}}\exp\left(-\frac{\left(\theta_{i'1}-\theta_{i'}\right)^2}{2\sigma_0^2}\right)$$

$$\propto \exp\left(-\frac{\sigma_0^2\left(\theta_{i'}-\mu\right)^2+\tau^2\left(\theta_{i'1}-\theta_{i'}\right)^2}{2\tau^2\sigma_0^2}\right)$$

$$\propto \exp\left(-\frac{\left(\theta_{i'}-\frac{\mu\sigma_0^2+\theta_{i'1}\tau^2}{\sigma_0^2+\tau^2}\right)^2}{\frac{2\tau^2\sigma_0^2}{\sigma_0^2+\tau^2}}\right)$$

$$\propto N\left(\frac{\mu\sigma_0^2+\theta_{i'1}\tau^2}{\sigma_0^2+\tau^2},\frac{1}{\frac{1}{\sigma_0^2}+\frac{1}{\tau^2}}\right)$$

### S1.3. Pólya-Gamma data augmentation

It is difficult to directly sample from the conditional posterior of $p\left(\theta_{ij}|X,\Theta\setminus\theta_{ij}\right)$ and $p(\theta_{i'1}|X,\Theta\setminus\theta_{i'1})$ as these are intractable.

$$p\left(\theta_{ij}\middle|X,\Theta\setminus\theta_{ij}\right)\propto\binom{n_{ij}}{x_{ij}}\frac{\left(e^{\theta_{ij}}\right)^{x_{ij}}}{\left(1+e^{\theta_{ij}}\right)^{n_{ij}}}\times(\sigma_i^2)^{-\frac{1}{2}}\exp\left(-\frac{\left(\theta_{ij}-\theta_i\right)^2}{2\sigma_i^2}\right)$$

Therefore, we instead implement a data augmentation strategy that is based on Pólya-Gamma (PG) latent variables[1]. We further define auxiliary variables $\lambda_{ij}$ and $\lambda_{i\prime 1}$, which, if conditional on $\theta_{ij}$ and $X$, follow the Pólya-Gamma distribution.

$$p\left(\lambda_{ij}|\theta_{ij},X\right)\sim PG\left(n_{ij},\theta_{ij}\right)$$

$$p(\lambda_{i\prime 1}|\theta_{i\prime 1},X)\sim\ PG(n_{i\prime 1},\theta_{i\prime 1})$$

Based on the Pólya-Gamma data augmentation strategy, let $\lambda_{ij}\sim PG\left(n_{ij},0\right)$, the following identity holds for all $x_{ij}>0$:

$$\frac{\left(e^{\theta_{ij}}\right)^{x_{ij}}}{\left(1+e^{\theta_{ij}}\right)^{n_{ij}}}=2^{-n_{ij}}e^{\kappa_{ij}\theta_{ij}}\int_0^\infty e^{-\frac{\lambda_{ij}\theta_{ij}^2}{2}}p\left(\lambda_{ij}\right)d\lambda_{ij}$$

Where $\kappa_{ij}=x_{ij}-n_{ij}/2$. The conditional distribution of $\lambda_{ij}$ given $\theta_{ij}$ and $X$ is given by

$$p\left(\lambda_{ij}\middle|\theta_{ij},X\right)=\frac{e^{-\frac{\lambda_{ij}\theta_{ij}^2}{2}}p\left(\lambda_{ij}\right)}{\int_0^\infty e^{-\frac{\lambda_{ij}\theta_{ij}^2}{2}}p\left(\lambda_{ij}\right)d\lambda_{ij}}$$

$$=\frac{e^{-\frac{\lambda_{ij}\theta_{ij}^2}{2}}p\left(\lambda_{ij}\right)}{E_{\lambda_{ij}}\left[\exp\left(-\frac{\lambda_{ij}\theta_{ij}^2}{2}\right)\right]}$$

$$\sim PG(n_{ij},\theta_{ij})$$

Therefore, we have

$$p\left(\theta_{ij}\middle|X,\Theta\setminus\theta_{ij},\lambda_{ij}\right)\propto e^{\kappa_{ij}\theta_{ij}}p(\lambda_{ij}|\theta_{ij},X)\times(\sigma_i^2)^{-\frac{1}{2}}\exp\left(-\frac{\left(\theta_{ij}-\theta_i\right)^2}{2\sigma_i^2}\right)$$

$$\propto\exp\left(\kappa_{ij}\theta_{ij}-\frac{\lambda_{ij}\theta_{ij}^2}{2}\right)(\sigma_i^2)^{-\frac{1}{2}}\exp\left(-\frac{\left(\theta_{ij}-\theta_i\right)^2}{2\sigma_i^2}\right)$$

$$\propto \exp\left(-\frac{\left(\theta_{ij}-\frac{\kappa_{ij}}{\lambda_{ij}}\right)^2}{2\frac{1}{\lambda_{ij}}}-\frac{\left(\theta_{ij}-\theta_i\right)^2}{2\sigma_i^2}\right)$$

$$\propto \exp\left(-\frac{\left(\theta_{ij}-V_{ij}\right)^2}{m_{ij}}\right) \sim N\left(V_{ij}, m_{ij}\right)$$

The conditional posterior based on the defined auxiliary parameters can thus be written as the form of known distribution

$$p\left(\theta_{ij} | X, \Theta \setminus \theta_{ij}, \lambda_{ij}\right) \quad \propto N\left(V_{ij}, m_{ij}\right)$$

where

$$V_{ij} = \left(\lambda_{ij} + \frac{1}{\sigma_i^2}\right)^{-1}$$

$$m_{ij} = V_{ij}\left(\kappa_{ij} + \frac{\theta_i}{\sigma_i^2}\right)$$

Thus, the Gibbs sampler of the BIT model is now only comprising of known distributions, with $\lambda_{ij}$ can be sampled from PG distribution[1].

$$p\left(\theta_{ij} \middle| X, \Theta \setminus \theta_{ij}, \lambda_{ij}\right) \sim N\left(V_{ij}, m_{ij}\right)$$

$$p\left(\lambda_{ij} \middle| \theta_{ij}, n_{ij}\right) \sim PG(n_{ij}, \theta_{ij})$$

### Supplementary Note 2.

#### Extended discussion of BIT identified TRs for each cancer type.

Breast cancer – The used TCGA cancer-type specific accessible regions were mostly derived from invasive breast carcinoma, which is a broad category of breast cancer that can spread from the original ducts or lobules into the surrounding breast tissue. BIT identified Top TRs include FOXA1, ESR1, TET2, NCOA3, PGR, AR, and GATA3. These TRs can formulate a complex network to regulate hormone-responsive genes, crucial for breast cancer cell proliferation and survival. FOXA1 facilitates estrogen receptor (ER, encoded by ESR1) binding to DNA, enhancing ER-mediated transcription[2]. The progesterone receptor (PR, encoded by PGR) further modulates this signaling, indicating effective ER activity[3]. GATA3 supports luminal cell differentiation and is often co-expressed with ER, contributing to the maintenance of a differentiated state[4]. The androgen receptor (AR) can interact with ER signaling pathways, influencing cancer cell growth in a context-dependent manner[5]. NCOA3 often acts as a coactivator and has strong inverse relationship with the expression of ER, promoting tumor growth[6]. TET2, involved in DNA demethylation, helps regulate gene expression epigenetically, potentially impacting the function of these transcription factors[7]. Given that hormone receptor-positive breast cancer constitutes a significant proportion of all breast cancer cases (~70-80%), it is unsurprised these TRs are ranked in top positions.

Bladder cancer - Urothelial carcinoma, also known as transitional cell carcinoma, is the most common type of bladder cancer. It originates in the urothelial cells lining the inside of the bladder. GRHL2 has been linked to epithelial-mesenchymal transition (EMT) [8], which is associated with cancer metastasis[9]. TP63 and TP73 are members of p53 transcription factor family, which may act as tumor suppressor by regulating cell cycle and apoptosis, previous studies reveal the different expression of isoforms in p63 can be related to the urothelial development and bladder carcinoma progression[10]. GATA3 is another TF worth to be noted. The loss of which can promote cell migration and invasion in bladder cancer[11]. ARID1A and

SMARCB1 are component of the SWI/SNF chromatin complex, which modulate the chromatin structure and potentially regulate gene expression in bladder cancer[12]. ARID1A has been shown to be related to the self-renewal of bladder cancer stem cells[13].

Colon cancer – Colon adenocarcinoma is the most common type of colon cancer, originating from the glandular epithelial cells lining the colon. It is a type of colorectal cancer (CRC), which encompasses cancers of the colon and rectum. In the BIT identified TRs, HNF4A and HNF4G are known to be important for the epithelial cell differentiation[14-16]. CDX2 is known tumor suppressor in colorectal cancer[17], loss of CDX2 can lead to poor prognosis and more aggressive tumor behavior. ETV5 and ELF3 are both ETS family transcription factors, evidence has associated the expression of two factors with the poor prognosis of CRC and shown the two TRs can contribute to the tumor progression and promote CRC angiogenesis[18,19].

Lung cancer – Epigenomics regions mostly derived from lung adenocarcinoma, the most common subtype of non-small cell lung cancer. NKX2-1 is one of the most well-known transcription factors in the lung adenocarcinoma[20], which is critical for the normal differentiation and development of lung epithelial cells, and commonly used as a diagnostic marker. FOSL2 and JUNB are all AP-1 transcription factor subunit. Recent studies claim the activity change of AP-1 transcription factor complex can be related to the cell transformation from normal to adenocarcinoma[21], potentially a reason why these TRs are ranked to top 10. SMAD3 can contribute to the progression of lung cancer through the TGF-ß signaling, the alternation of which has been linked to cancer progression[22,23]. Other TRs can have more general roles in lung cancer. For instance, FOXA2 is reported to be a suppressor of tumor metastasis by inhibition of EMT in human lung cancers[24].

Liver cancer – Liver hepatocellular carcinoma (HCC) is the most common type of primary liver cancer. It originates from hepatocytes, the main type of liver cell, and is a major cause of cancer-related mortality worldwide. BIT identified TRs such as HNF4A and HLF are known for regulating liver-specific genes and maintaining hepatocyte differentiation, dysregulation of these TRs can be associated to the progression of liver cancer[25,26]. Another TR worth to be noted is NR1H4, also known as the Farnesoid X Receptor (FXR). FXR acts as a tumor suppressor by regulating bile acid homeostasis and protecting against liver inflammation and fibrosis, which can lead to HCC. Loss of FXR function is associated with liver carcinogenesis[27].

Mesothelium cancer – Regions were derived from mesothelioma, a rare and aggressive cancer that develops from the mesothelial cells lining the pleura (the protective lining of the lungs) and other serosal surfaces. Unlike other cancer types, there are less studies of the function of transcriptional regulators in mesothelioma. TRs that are worth to be noted include PPAR$\gamma$, TCF21, and TWIST1. PPAR$\gamma$ activation is associated with plural mesothelioma invasion[28]. TCF21 and TWIST1 can be indirectly associated with mesothelioma by promoting EMT, which is associated with increased invasiveness and metastasis in mesothelioma[29].

Prostate cancer – prostate adenocarcinoma is one of the most common cancers in males and a leading cause of cancer-related deaths, which originates in the glandular cells of the prostate, a small gland that produces seminal fluid in males. For the identified TRs, dysregulation of Androgen receptor (AR) signaling is crucial for the growth and survival of prostate cancer cells[30]. PIAS1 is a co-regulator of AR[31], and FOXA1 is a pioneer factor helps AR access target gene promoters[32]. HOXB13 and NKX3-1 are also associated with increased prostate cancer risk[33,34].

Squamous cancer – Squamous cell carcinoma (SCC) is a malignancy that arises from squamous epithelial cells. It can occur in any organ that contains squamous epithelium, including the skin, lungs, esophagus, cervix, and head and neck region. Input epigenomic regions are derived from multiple squamous cell cancer types including head and neck squamous cell carcinoma, cervical squamous cell carcinoma, lung squamous cell carcinoma, and esophageal carcinoma. Multiple BIT identified TRs have strong association with squamous cell carcinoma, such as p53 family TRs TP53, TP63, and TP73. Mutation of which can promote the tumor progression in squamous cells[35-37]. MAML1 is a coactivator in notch signaling, which is often dysregulated in SCC[38]. Other TRs including KLF5 and SOX2 are often over- or under-expressed in SCC and can promote tumorigenesis by enhancing cell proliferation and survival[39,40].

Testicular cancer – Testicular germ cell tumors (TGCTs) are cancers that originate from the germ cells in the testes. They are the most common malignancy in young men aged 15-35 years. TGCTs are classified into two main subtypes: seminomas and non-seminomas. The function of NANOG[41], SOX17[42], SOX2[43], TFAP2C[44], and POU5F1[45] have been established in the previous studies for the germ cells development and differentiation. The mutation of which can be associated with the development of testicular cancer.
